# Integral Synthesis and Clearance Analysis via DIA (ISDia) Reveals Diverse Protein Abundance Regulation during Endoplasmic Reticulum Stress

**DOI:** 10.1101/2025.07.23.666381

**Authors:** Yue Dou, Danny Qiu, Vivien Li, Maja E. Wierzbińska, Gregory R. Keele, Wenpeng Liu, Jie Yang, Joao A. Paulo, Ling Qi, Tian Zhang

## Abstract

Endoplasmic Reticulum (ER) stress disrupts protein homeostasis, driving cellular responses critical for survival, development and disease. However, systematic proteome-wide analyses of protein synthesis and clearance during ER stress remain limited. In this study, we developed Integral Synthesis and clearance analysis via DIA (ISDia), a robust mass spectrometry (MS)-based workflow that integrates multiple pulsed-SILAC labeling timepoints under one condition into a single sample and utilize data-independent acquisition (DIA) to quantify heavy and light peptide changes across conditions. It enables proteome-wide tracking of protein synthesis and clearance with high coverage by directly comparing light and heavy peptides in pSILAC-DIA experiments across multiple conditions. Using ISDia, we uncover diverse regulatory mechanisms by which protein synthesis and clearance are modulated to regulate protein abundances during ER stress, revealing PERK dependent and independent regulatory mechanisms across subcellular compartments, complexes and isoforms. These findings highlight the potential of ISDia as a powerful and widely applicable workflow for revealing protein abundance regulatory mechanisms.

## INTRODUCTION

Protein homeostasis relies on the tightly coordinated regulation of protein synthesis, protein folding and protein clearance. Disruption of this balance is a common driver of cellular stress across diverse diseases, including metabolic syndromes, cancer, immune disorders, and neurodegenerative conditions^1^. The endoplasmic reticulum (ER) plays a key role in proteostasis. The accumulation of misfolded proteins in the ER activates the unfolded protein response (UPR) to alleviate ER stress by attenuating translation, increasing chaperone capacity, and promoting ER-associated degradation (ERAD)^2–4^. During the UPR, the PERK branch represent a major mechanism for attenuating global protein translation through eIF2α phosphorylation and activating autophagy by ATF4^2,5^. Prior proteomic profiling has provided broad views of ER stress-induced changes in total protein abundance and turnover, but these measurements cannot disentangle the respective regulation of protein synthesis and clearance to overall protein abundance changes^6,7^. The proteome-wide consequences of PERK signaling for protein synthesis and clearance remain incompletely understood. This gap has highlighted the need for proteomic analysis that can directly capture changes in protein synthesis and clearance under ER stress.

Pulsed Stable Isotope Labeling by Amino Acids in Cell Culture (pSILAC) coupled with MS-based proteomics provides a way to directly track steady-state protein turnover, *i.e*, the continuous renewal of proteins through protein synthesis and clearance^8–11^. In a standard pSILAC, cells are switched to medium containing heavy isotope labeled amino acids, which are incorporated into newly synthesized proteins, whereas pre-existing proteins remain light-labeled. Consequently, accumulation of heavy labeled peptides reflects protein synthesis, while loss of the light labeled pool reflects protein clearance (i.e., the sum of protein degradation and dilution effect of cell division). Under steady-state conditions, synthesis, degradation and turnover rates can be calculated by fitting heavy to light peptide (H/L) ratios at different time points based on first-order kinetics^12,13^. However, under many biological processes, including ER stress, cells often deviate from steady-state conditions, such that protein synthesis and clearance flux are no longer equal. The steady-state assumptions underlying kinetic models may not hold under these conditions. Protein synthesis and degradation should therefore be analyzed separately.

Recenlty, an increasing number of studies have moved beyond bulk turnover estimation to quantify protein synthesis and degradation separately using pSILAC. Boisvert *et al.* used a three-channel pSILAC strategy to measure protein synthesis, degradation, and turnover rates under steady states^14^. Fierro-Monti *et al.* later developed a pulse-chase variant of SILAC, enabling relative quantitation of newly synthesized and pre-existing protein level comparison under two conditions to examine protein synthesis and degradation^15^. Because pSILAC is limited by the number of isotopic labeling channels, the number of experimental conditions that can be incorporated within a single experimental design is restricted. More recently, studies has intergrated pSILAC with tandem mass tag (TMT) labeling, allowing for multiplexed analysis of protein synthesis and degradation by tracking newly synthesized and pre-existing proteins under different time points^13,16^ or conditions^17^. In particular, Mutiplexed Proteome Dynamics Profiling (mPDP) enabled protein degradation to be investigated without relying on kinetic equations derived under steady-state assumption ^17^. However, TMT-SILAC workflows require additional labeling and often extensive fractionation. Moreover, because heavy-and light-labeled peptides appear as distinct precursor ions, stochastic precursor selection in DDA may result in only one form being fragmented, reducing the completeness of paired heavy and light quantification. ^13,18–20^.

Recent advances in data-independent acquisition (DIA) approach has enabled deeper proteome coverage and robust quantitation with simplified workflow^21^. Previously, pSILAC has been coupled with DIA to quantify protein turnover^18,22–25^ or to examine protein synthesis by enriching peptides labeled with clickable methionine analogs under different conditions^26^. However, pSILAC-DIA has not been systematically explored to simultaneously investigate the contribution of protein synthesis and clearance in regulating protein abundance. Here, we leveraged high-sensitivity MS instrumentation and demonstrated that heavy and light intensities in pSILAC-DIA can be directly compared across samples, thereby establishing a foundation for directly comparing protein synthesis and clearance across conditions. By integrating heavy and light intentities with protein total abundance, we are able to distinguish the contribution of protein syntheis and degradation to protein abundance changes. Based on this, we developed ISDia (Integral Synthesis and clearance analysis via DIA), which pools multiple pSILAC labeling time points into a single mixture to increase proteome coverage and quantitation robustness, providing a streamline workflow to examine protein synthesis and clearance regulation across conditions.

We then applied ISDia to investigate protein abundance regulation during ER stress, revealing that cells displayed diverse synthesis and clearance regulatory mechanisms to regulate protein abundance across individual proteins, subcellular compartments and protein complexes. To delineate the PERK-dependent and independent synthesis regulation during the UPR, we applied ISDia to PERK^-/-^ cells and found that mitochondrial protein synthesis is selectively preserved during the global translation suppression in a PERK-dependent manner. Moreover, protein complex analysis revealed PERK-mediated isoform-specific regulation of p97/VCP during the UPR. Altogether, our study presents a streamlined proteomics workflow that uncovers mechanisms underlying proteome remodeling during the UPR by quantifying the changes of protein synthesis and clearance.

## RESULTS

### DIA enables accurate and reproducible cross-sample quantitation of heavy and light peptides

To establish ISDia, we tested whether heavy and light intensities measured by DIA on an Orbitrap Astral MS could track abundance changes in newly synthesized and pre-existing proteins across samples in pSILAC experiments. We generated standards with predefined H/L ratios by mixing light-labeled cells and fully heavy-labeled cells at H/L ratios of 1:27, 1:9, 1:3, 1:1, 3:1, 9:1, and 27:1 to model the relative proportions of newly synthesized and pre-existing proteins under different conditions (**Fig. 1A, S1A**). Equal amounts of peptides were injected across all runs. All DIA files were analyzed in Spectronaut in a single search workflow, with cross-run normalization enabled to correct run-to-run technical variation. We quantified both heavy and light intensities for more than 10,000 proteins at each mixing ratio in a single run, with 8,088 to 10,726 proteins quantified across three replicates per mixing ratio (**Fig. 1B**). We calculated coefficients of variation (CVs) across three replicates for light, heavy, total (sum of heavy and light, H+L) protein intensity, and the H/L ratio at each mixing ratio. Proteins were considered reproducibly quantified if their CV was ≤ 20% across replicates. Across the mixing ratios, 4,456 to 8,455 proteins met this threshold for all four quantitative measurements. We observed that the CVs of total intensity were below 20%, indicating high reproducibility, whereas higher CVs were observed when the corresponding heavy or light channel was present at low abundance (**Fig. 1C**). As a result, H/L ratio measurements showed greater variability at extreme H/L ratios (**Fig. 1C**). Despite the high CV at certain measurements, the expected H/L ratios were accurately recapitulated in most standards (**Fig. 1D**). These results demonstrated robust quantitation in pSILAC-DIA samples and highlighted the challenges of accurately quantifying low abundance proteins in pSILAC samples.

**Figure 1.**
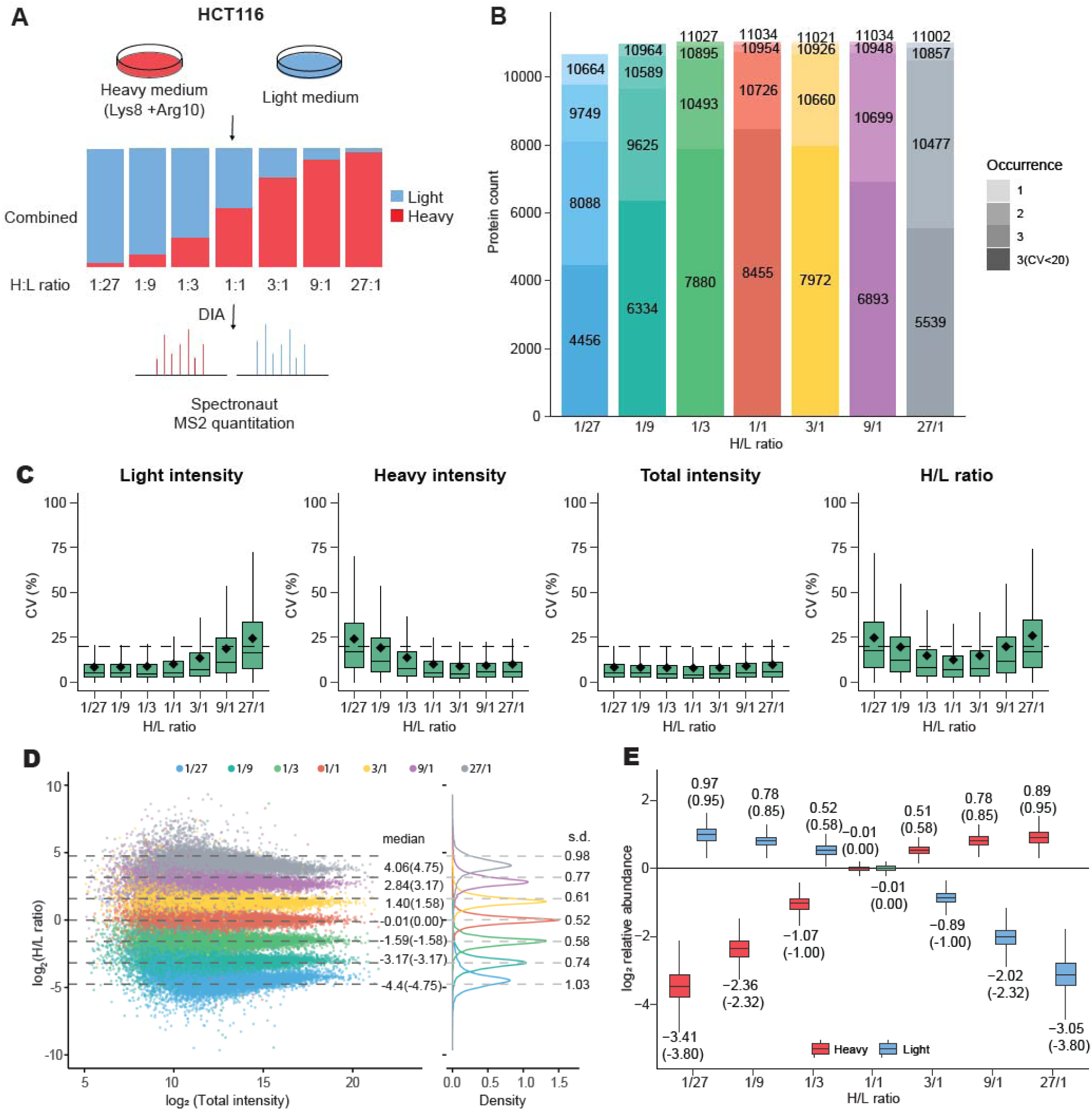
Predefined H/L ratio mixtures benchmark pSILAC-DIA for accurate within and across-sample quantitation. A. Workflow for generating standards with pre-defined H/L ratios. B. Proteome coverage achieved across standards. Each bar is labeled with four numbers representing the number of proteins for which both light and heavy intensities were detected in one, two, or all three replicates, and the number consistently quantified across all three replicates with a coefficient of variation (CV) < 20% for light, heavy, and total intensities as well as H/L ratios. C. Boxplots showing the CV of light, heavy, total peptide intensities and H/L ratios across three replicates of the standards. The black dashed line indicates the 20% CV threshold. D. Comparison of protein-level H/L ratio distribution in standards. s.d. indicates the standard deviation of log_₂_(H/L) ratios within each group; Dashed lines mark theoretical log_₂_(H/L) values: log_2_(1/27)=-4.75, log_2_(1/9)=-3.17, log_2_(1/3)=-1.58, log_₂_(1) = 0, log_₂_(3) = 1.58, log_₂_(9) = 3.17, and log_₂_(27) = 4.75. E. DIA-based cross-sample quantitation of light and heavy peptide intensities in standards. Light and heavy intensities were normalized to those from the 1:1 standard sample. Theoretical log_2_-transformed fold changes were labeled in parentheses.

We then evaluated quantitation accuracy across samples by normalizing heavy and light intensities to their corresponding values in the 1:1 H/L standard, which served as the reference. We found that DIA measurements recapitulated the expected heavy and light peptide intensity differences across samples (**Fig. 1E**). To evaluate whether this observation was specific to the Orbitrap Astral MS, we searched and analyzed the DIA standards acquired on a Q-Exactive HF-X tandem MS^23^. We found that, except for extreme ratios such as 0.1/99.9 and 1/99, the measured heavy and light intensities were consistent with the expected difference across samples and showed high robustness with lower proteome coverage (**Fig. S1B-D**). Together, these findings demonstrate that light and heavy peptide intensity difference can be accurately quantified for cross-sample comparison in pSILAC-DIA experiments. This capability enables direct comparison of newly synthesized and pre-existing protein pool changes across conditions, thereby laying foundation of interpreting changes in protein synthesis and clearance across conditions. Furthermore, this approach overcomes the limited multiplexing capacity of pSILAC labeling channels, allowing quantitative comparisons across multiple samples without the need for additional isobaric labeling.

### Time-resolved pSILAC-DIA enables dissection of protein synthesis and clearance contributions to proteome dynamics during UPR activation

We next applied direct cross-sample intensity comparisons to track synthesis and clearance dynamics during UPR activation. In this experiment, we treated Hela cells with the N-glycosylation inhibitor tunicamycin (Tm) or the SERCA inhibitor thapsigargin (Th), two established ER stressors that activate convergent UPR signaling pathways. Activation of the UPR was confirmed by induction of key pathway components and downstream targets. Under both treatments, IRE1α-mediated splicing of X-box binding protein 1 (XBP1) mRNA and accumulation of the spliced XBP1s protein were detected by Western blot at 6h (**Fig. 2A**). For this proof-of-principle application, isotope labeling was initiated concurrently with DMSO, Tm, or Th treatment, and cells were harvested at 6, 12, 24, and 48 h post-treatment in biological triplicates (**Fig. 2B**). Since cells were still dividing during labeling, changes in light peptide intensity capture the combined contributions of protein degradation and dilution-driven clearance. Notably, as pSILAC labeling last for 48h, heavy peptide intensity is affected not only by protein synthesis but also by protein clearance. Therefore, interpretation of protein synthesis changes requires consideration of both heavy intensity alteration and protein clearance that reflected by light peptide intensity. This time-resolved design enabled separate tracking of newly synthesized and pre-existing protein pools across the stress response, allowing us to examine how changes in synthesis and clearance contribute to changes in total protein abundance over time.

**Figure 2.**
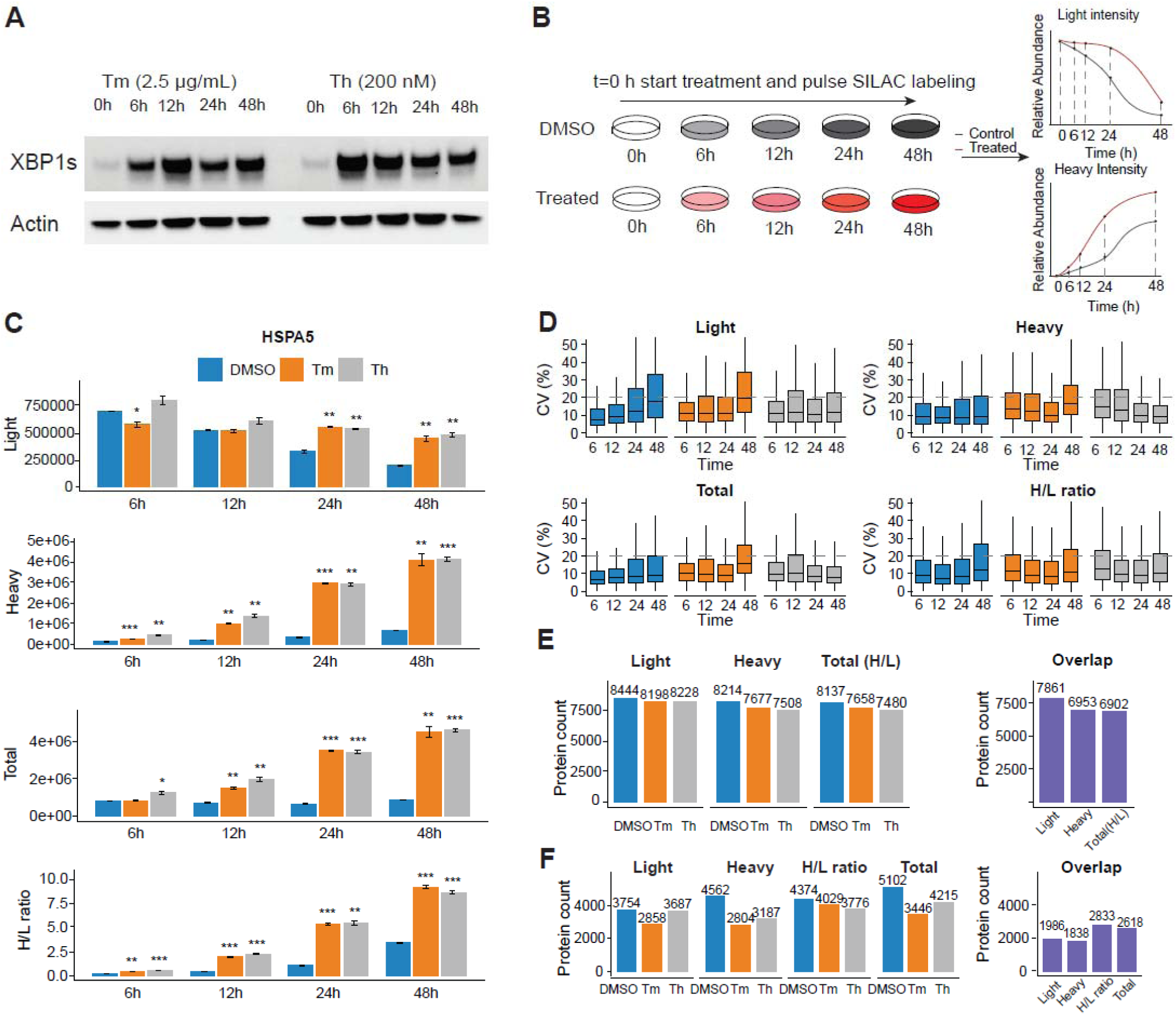
Time-resolved pSILAC-DIA analysis of protein synthesis and clearance during the UPR. A. Western blotting analysis of XBP1 protein levels in cells treated with 2.5µg/mL tunicamycin (Tm) or 200mM thapsigargin (Th) for 0, 6, 12, 24 and 48 hours. B. Schematic of pSILAC labeling workflow used for the time-resolved pSILAC-DIA experiment. C. Bar plots showing light, heavy, H+L intensity and H/L ratio measurements for HSPA5 in HeLa cells treated with DMSO, Tm, or Th for 6, 12, 24, and 48 h. Values represent mean ± SEM of biological triplicates. Statistical significance is indicated as * p < 0.05, ** p < 0.01, *** p < 0.001. D. Boxplots showing the CVs for light, heavy, total (H+L) peptide intensities and H/L ratios across replicates. The dashed line indicates the 20% CV threshold. Only proteins were quantified in three replicates were included for CV calculation. E. Number of proteins quantified across three biological replicates at all four time points for each condition. Bar plots show the numbers of proteins with quantified light intensities, heavy intensities, and total (H/L) under each condition. The right panel shows the overlapping proteins in three conditions for each metric. Total and H/L ratio values are calculated only for replicates with both light and heavy intensities quantified. F. Number of proteins quantified across three biological replicates at all four time points for each condition after CV < 20% filtering.

We analyzed heavy and light intensity changes at each timepoint under DMSO, Tm and Th treatment and showed HSPA5 as an illustrative example (**Fig. 2C, Table S1**). HSPA5, also known as ER chaperon BiP, operates as a Hsp70 chaperon in complex with its cochaperone proteins. It serves as a sensor of ER stress, and its expression gets upregulated under UPR^27–29^. Interestingly, heavy peptide intensities increased more rapidly under ER stress than in DMSO beginning at 6h. (**Fig. 2C**). Light peptide intensities of HSPA5 did not show a significant increase until 24h and later time points, reflecting that the increased total HSPA5 abundance were driven primarily by increased protein synthesis under ER stress at the early time-points. After 24h, both increased protein synthesis and reduced protein clearance contribute to the increase in total HSPA5 abundance under both treatments. Collectively, these results demonstrated dynamic protein synthesis and clearance regulation during UPR activation.

We next evaluated the quantitative robustness and proteome coverage of these time-resolved pSILAC-DIA samples. Consistent with the trend observed in the standards, heavy and light intensities were robustly quantified except at time points where the corresponding channel was present at relatively low abundance (**Fig. 2D**). Across the four time points analyzed, we quantified light, heavy, and total intensities, as well as H/L ratios, for approximately 8,000 proteins under each condition, with about 7,000 proteins overlapping across the three conditions (**Fig. 2E**). However, applying a 20% CV threshold substantially reduced the number of proteins that were reproducibly quantified across all four time points and shared across all three conditions (**Fig. 2F**). This reduction was likely due to both lower quantitative reproducibility when heavy or light intensities were low and the stringent requirement for robust quantification across all four time points (**Fig. 2D**). To test it, we then examined the proteome coverage and quantitative reproducibility at 48h time point. At 48h, we quantified around 7,800 proteins under each condition (**Fig. S2A**). Applying a CV threshold of <20% markedly reduced the number of proteins with robust measurements across all four matrices (**Fig. S2B**). Together, these results highlighted the challenge of reproducibly quantifying low abundance peptide and the need for a more robust workflow for quantifying protein synthesis and clearance with higher reproducibility and deeper proteome coverage.

### Development of the ISDia method to investigate protein synthesis and clearance regulation during ER stress

To increase the throughput and improve quantitative coverage and robustness, we pooled equal amounts of protein from each time point within same condition in pSILAC-DIA experiments before MS analysis. We then applied the same search method as described above and quantified changes in protein synthesis, clearance and protein abundance by directly comparing integrated light, heavy intensities and H+L across conditions. We term this method Integral Synthesis and clearance analysis via DIA (ISDia) (**Fig. 3A**).

**Figure 3.**
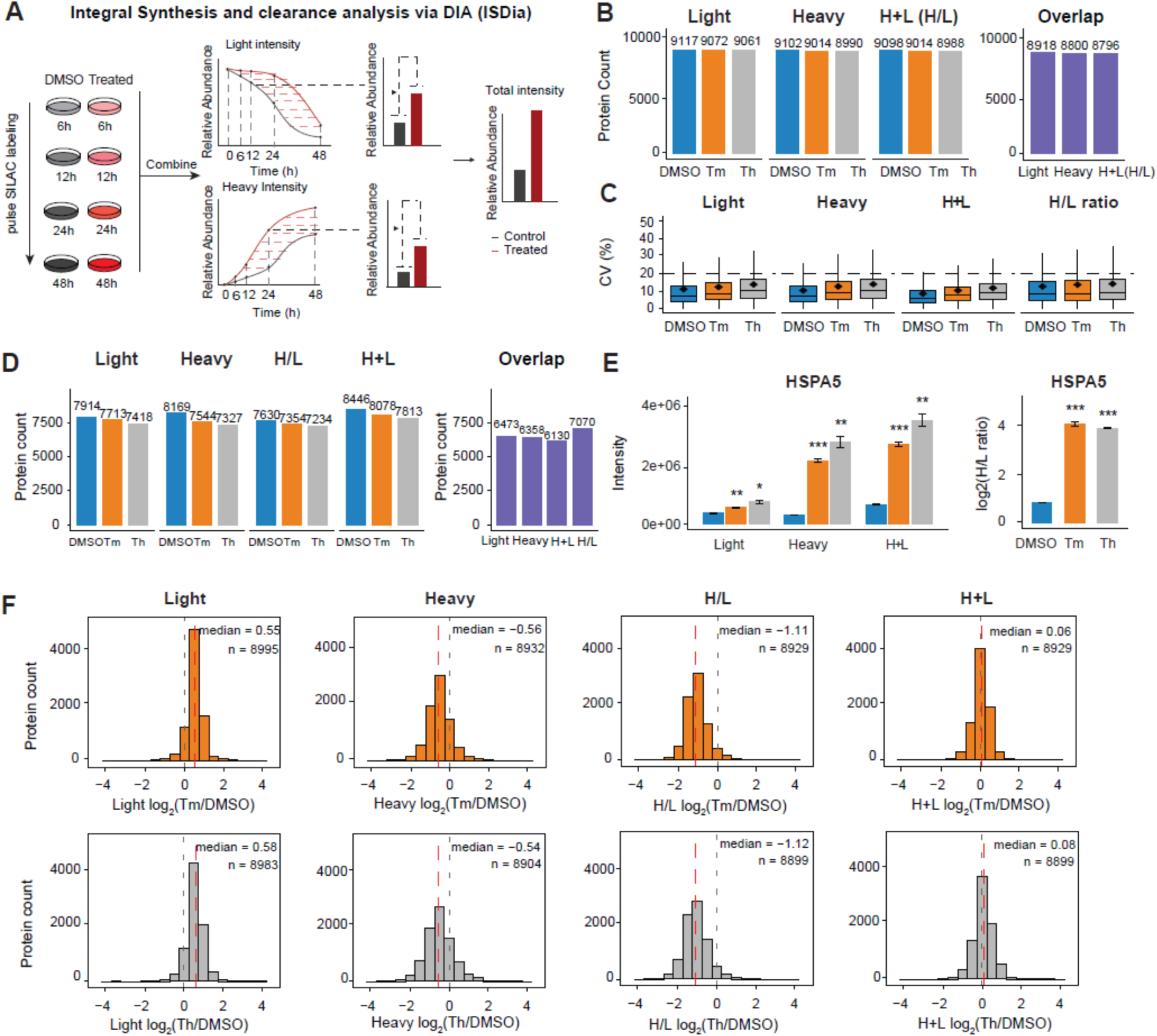
ISDia enabled increased proteome coverage and quantitative reproducibility. A. Schematic of the ISDia workflow. B. Number of proteins quantified by ISDia with light, heavy, and H+L intensities (H/L ratio) across three replicates. H+L and H/L ratio values were calculated only when both light and heavy intensities were quantified. C. Boxplots showing the CVs of light, heavy, H+L intensities, and H/L ratios across three biological replicates for different conditions. The black dashed line indicates the 20% CV threshold used for data quality filtering. Only proteins with heavy and light intensities quantified in three replicates were included for CV calculation. D. Number of proteins quantified by ISDia with light, heavy, and H+L intensities (H/L ratio) across three replicates after applying CV < 20% filtering. H+L and H/L ratio values were calculated only when both light and heavy intensities were quantified. E. Changes in light, heavy, and H+L intensities, as well as H/L ratio, for HSPA5 following Th or Tm treatment relative to DMSO. Statistical significance: ***□p□<□0.001; **□0.001□<□p□<□0.01; *□0.01□<□p□<□0.05. F. Distribution of log_₂_ fold changes for light, heavy, H+L intensities and H/L ratio, under Tm (top) and Th (bottom) treatment relative to DMSO control. H+L intensity and H/L ratio values were calculated only when both light and heavy intensity measurements were available. *n* indicates the number of proteins included in each panel.

To evaluate how pooling scheme affects proteome coverage and quantitative reproducibility we performed pSILAC labeling across 8 time points (4, 8, 12, 24, 36, 48, 60, and 72 h) in HeLa and RPE1 cells under steady state independently. We then compared pooling schemes varying in both the number of time points included (2, 4, or 8) and their position within the time course (early, middle, or late), evaluating effects on proteome coverage and replicate variability **(Fig. S3A)**. Across both cell lines and both channels, we observed two consistent patterns. First, proteome coverage increased as additional time points were incorporated into the pooled sample, with eight-time-point pooling providing greater coverage than four or two time point pooling (**Fig. S3B**). This improvement is likely due to pooling reduced dependence on any single time point with low-abundance peptides, thereby increasing the likelihood of peptide identification. Second, among schemes containing the same number of time points, coverage and quantitation precision decreased when the pooling window was shifted toward later time points (**Fig.S3B-C**). This trend is expected for the light channel because, with prolonged labeling, light proteins are progressively depleted through degradation and cellular dilution, resulting in lower peptide intensities, reduced detectability and poorer quantitation precision.

We applied ISDia to characterize protein clearance and synthesis alteration under ER stress conditions by pooling equal amounts of protein from all four sampled time points (6,12,24,48 h). We quantified light, heavy and calculated their sums and ratios for approximately 9,000 proteins under each condition (**Fig. 3B, Table S2**). CVs across replicates were below 20% for the majority of the quantified proteins (**Fig. 3C**). Using a 20% CV threshold as a reproducibility criterion, ∼7,500 proteins were quantified per metric (light, heavy, H+L or H/L ratio), with ∼6,500 proteins consistently quantified across all three conditions. These results demonstrated higher proteome coverage and robust quantitation compared to time resolved pSILAC-DIA (**Fig. 2D-F**, **Fig. 3C-D**). In addition, ISDia reduced the analytical burden by decreasing the number of samples required for analysis from 36 to 9.

To understand the biological meaning of the ISDia fold change, we mathematically modeled how pooling equal amounts of proteins across time points determines the resulting pooled fold change (**Method**). This analysis showed that the fold change in ISDia is jointly determined by two factors: the treatment-to-control fold change at each time point (*FC_i_*) and the relative contribution of that time point to the pooled control signal (*w_i_*). Accordingly, it represents an abundance-weighted average response, in which the contribution of each time point is influenced by peptide abundance, labeling kinetics, and the magnitude of the biological response. We then examined how individual time points contributed to the pooled fold changes measured by ISDia under ER stress. For each time point, we quantified both its relative contribution to the pooled signal and its corresponding treatment-to-control fold change. Consistent with the kinetics of pSILAC labeling, later time points contributed to an increasing proportion of the pooled heavy-channel signal, reflecting the accumulation of newly synthesized proteins, whereas their contribution to the light-channel signal decreased as pre-existing proteins were progressively cleared (**Fig. S3D**). The magnitude of the treatment-to-control fold change in the heavy channel remained relatively consistent across the time course, while the treatment effect in the light channel became progressively larger at later time points under ER stress (**Fig. S3E**). As an example, ISDia recapitulated the integrated change of HSPA5 in the time-resolved pSILAC-DIA experiment (**Fig. 2C, 3E**).

We next tested whether the responses identified in time-resolved experiments are preserved after pooling of samples in ISDia. Proteins were classified as exhibiting sustained, transient, or biphasic responses based on their treatment-induced changes at 6, 12, 24, and 48 h. Representative proteins illustrating each temporal response pattern are shown in **Figure S4A**. We assessed whether ISDia retained the same significant direction of change for sustained and transient responses (**Fig.S4B**). For proteins exhibiting sustained responses, the ISDia pooling strategy recapitulated approximately 90% of the responses identified by heavy-channel intensity and the H/L ratio, whereas approximately 60% of those identified by light-channel and total intensities were retained. We found that responses that were not retained generally exhibited smaller fold changes in ISDia (**Fig. S4D-F**). The light channels retained less responses, likely due to the temporal mismatch between effect size and signal contribution (**Fig. S3D-E**). Specifically, stronger treatment effects in the light channel occurred when its contribution to the pooled signal was reduced. As a result, changes in light-channel intensity were diluted and attenuated during pooling, which also influenced the total (H+L) measurement. For proteins with transient responses, ISDia captured approximately 50% of the changes observed in heavy intensity and the H/L ratio. Compared with sustained responses, temporally restricted changes were less frequently retained, indicating that transient signals are more susceptible to dilution during pooling (**Fig. S4B**). For biphasic responses, we assessed whether ISDia masked or captured one direction response. As expected, most of the biphasic-response proteins did not exhibit significant changes in ISDia (**Fig.S4C**), suggesting that opposing regulation effects may have canceled each other out, whereas a subset of proteins showed a significant change in one direction. In all, ISDia provides an integrated readout of protein regulation on protein clearance and synthesis over the selected labeling window. It is well suited for capturing monotonic responses, where sustained changes are more readily detected than transient ones.

We then performed global analysis to examine the responses under ER stress induced by Th and Tm treatments in ISDia. Samples from each condition were clustered separately in principle component analysis (**Fig. S5A**), Proteins showed increased light intensities, decreased heavy intensities and H/L ratios (**Fig. 3F**). These patterns suggest a global reduction in both protein synthesis and clearance. After cross-run normalization, the distribution of changes in total peptide intensity between ER stress and control conditions were centered around zero, while a subset of proteins exhibited substantial deviations, consistent with protein-specific abundance changes in response to treatment.

Multiple studies have characterized transcriptional and translational responses during the UPR using ribosome sequencing and proteomics technologies^30–35^ (**Table. S3**). To evaluate the concordance between ISDia and independent measurements of the UPR, we compared the results from ISDia and these external datasets. We first quantified the overlapping genes/proteins between ISDia and each external dataset and performed correlation analysis using the best matched parameters in each study (**Fig. S5B, Table. S3**). Although the overall correlations were moderate, likely reflecting differences in cell type, treatment duration, experimental design, and the distinct biological processes captured by each platform, ISDia recapitulated a substantial proportion of the strongest changes observed in the external datasets (**Fig. S5B**). Specifically, among gene products ranked within the top 5% of upregulated proteins or the bottom 5% of downregulated gene products in the external datasets, up to 90% showed concordant directional changes in ISDia (**Fig. S5B**). Overall, these analyses support that ISDia captures biologically consistent translation- and turnover-related responses in the most comparable external datasets.

In addition, we observed that Th- and Tm-induced changes in heavy and light peptide intensities, their sum, and H/L ratios were highly correlated, respectively, in the ISDia dataset (**Fig. S5C**). Given this strong concordance, we focused the detailed downstream analyses on the Th versus DMSO comparison as a representative ER stress condition and presented the relative result of Tm vs DMSO analysis in Supplementary **Figure S12-14**.

### Diverse regulation of protein synthesis and clearance modulated proteome changes during ER stress

To further investigate the regulatory mechanisms of protein abundance during ER stress, we color-coded individual proteins based on corresponding heavy, light and H+L intensity changes under treatment conditions and grouped them accordingly (**Fig. 4A-B**). Proteins exhibiting an absolute fold change of at least 1.5 and a p-value below 0.05 were considered significantly altered. The heatmap summarizes the changes in each protein group based on the median change across all four matrices (**Fig. 4C**), demonstrating that protein total abundance changes may originate from different combinations of protein synthesis and clearance regulation. In this experiment, heavy peptide intensity is affected by both protein synthesis and protein clearance. Therefore, we considered both heavy intensity and protein clearance alteration to interpret protein synthesis changes. We first focused on proteins with unchanged light intensities*, i.e.*, unchanged clearance. Among these, proteins in the blue group showed significantly increased heavy and total intensities. Coupled with the unchanged clearance, this result indicated that protein synthesis increased under Th treatment and drove the total protein abundance increase. We performed gene ontologies (GO) enrichment analysis for proteins in this group and found that they are involved in ER stress-related protein trafficking and quality control **(Fig. 4D)**. This observation is consistent with previous studies demonstrating these pathways were upregulated transcriptionally^36–38^. In contrast, proteins in the purple group showed significantly decreased heavy and total intensities, indicating reduced protein synthesis led to the decrease of total protein abundance. These proteins were enriched in cell cycle-related processes, including chromosome segregation, nuclear division, and mitotic spindle assembly checkpoint signaling **(Fig. 4E)**. Notably, many of these genes have been reported to be transcriptionally repressed during ER stress^39,40^. To distinguish changes in protein degradation from reduced protein dilution caused by slower cell division during the UPR, we derived cell-division offsets based on experimentally measured proliferation rates under each condition (**Methods, Fig. S6A-B**). For WT cells treated with Th, reduced cell division was predicted to shift a light-channel log_2_ fold change by +0.37, even if protein degradation remained unchanged. We therefore shifted the cutoff used to define significant changes by the cell-division offset, resulting in an adjusted interval of −0.21 to +0.95. Proteins outside this interval were interpreted as having significantly altered protein degradation. We found that a subset of the proteins in the purple groups fell below the lower cutoff (log_2_FC = -0.21), indicating significantly increased degradation under Th-treated conditions **(Fig. 4B)**. In contrast, most proteins in the blue group showed no significant change in degradation. This analysis provided an approximate estimate of treatment-induced changes in protein degradation.

**Figure 4.**
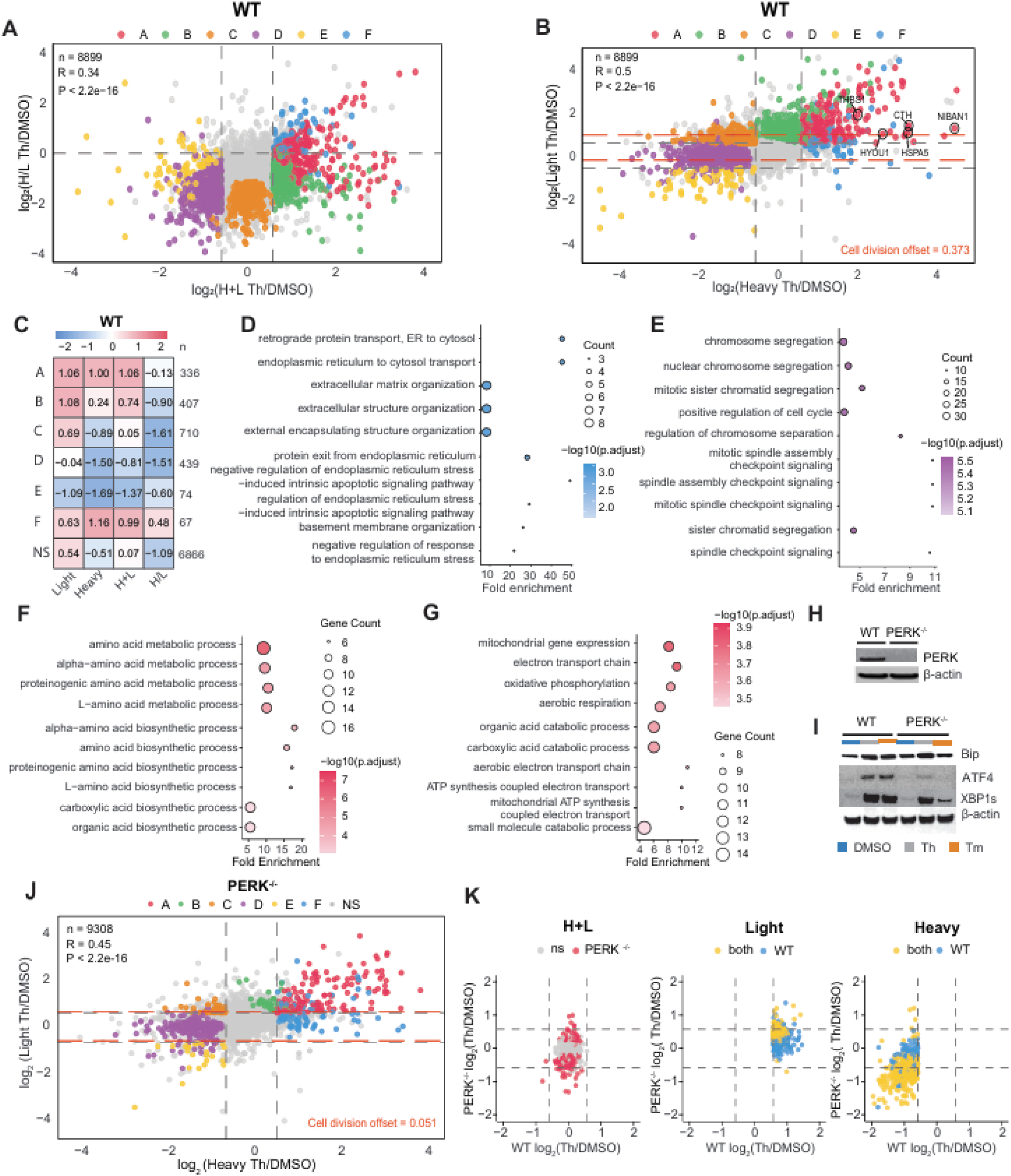
Diverse protein synthesis and clearance regulation shapes proteome abundance during the UPR. A. Scatter plot showing changes in H+L and H/L ratio between Th and DMSO treatments. Each point represents a protein. Grey dashed lines indicate the fold change thresholds of log_₂_(1.5) ≈ ±0.58 for H+L intensity, and log_₂_(1) = 0 for H/L ratio. Proteins are color-coded based on their changes across light, heavy, and H+L as follows: A: dark pink (up in light, heavy, and H+L); B: green (up in light and H+L, heavy not significant); C: orange (light up, heavy down, H+L not significant); D: purple (down in heavy and H+L, light not significant); E: yellow (down in light, heavy, and H+L); F: blue (up in heavy and H+L, light not significant) grey (others). Up is log_2_FC > 0.58 & p-value < 0.05; Down is log_2_FC < -0.58 & p-value < 0.05; no significant is p-value >0.05. n is the number of proteins shown; R is the Pearson correlation coefficient; p is the corresponding p-value. B. Scatter plot showing changes in heavy and light intensities between Th and DMSO treatment. Color scheme and significance thresholds are the same as in **A**. Red dashed lines indicate the cell division-adjusted light-channel fold-change thresholds, calculated as −0.58 + the cell division offset and +0.58 + the cell division offset. Cell division offset in WT is 0.373. C. Heatmap showing the median log_2_ fold change in light, heavy, H+L intensity, and H/L ratio for all proteins within groups highlighted in A and B. Values in each cell represent the group-wise median. N indicates the count of proteins in each group. D. Biological processes enriched among proteins in group F (blue). E. Biological processes enriched among proteins in group D (purple). F. Biological processes enriched among the proteins in group A (dark pink) with log_2_ H/L(FC) > 0 under Th treatment compared to DMSO. G. Biological processes enriched among the proteins in group A (dark pink) with log_2_ H/L(FC) < 0 under Th treatment compared to DMSO. H. Western blot analysis of PERK expression in untreated WT and PERK^-/-^ cells. I. Western blot analysis of downstream pathway markers in WT and PERK^-/-^ cells treated with 2.5 μg/mL tunicamycin (Tm) or 200 nM thapsigargin (Th) for 6 h. J. Scatter plot showing the changes in heavy and light intensities between Th and DMSO treatment in PERK^-/-^ cells. Color scheme and significance thresholds are the same as A. Red dashed lines indicate the cell division-adjusted light-channel fold-change thresholds, calculated as −0.58 + the cell division offset and +0.58 + the cell division offset. Cell division offset in PERK^-/-^ is 0.051. K. Scatter plot showing the log_2_(fold changes) in heavy, light and H+L intensities under Th treatment relative to DMSO between WT and PERK^-/-^ cells for group C (orange group) proteins filtered from WT cells (light up, heavy down, H+L not significant). Each point represents one protein from this defined group in WT, plotted by its corresponding log_2_(fold change) under Th relative to DMSO in WT and PERK^-/-^ cells. Colors represent significance of the displayed metric: gray, non-significant in both cell types; pink, significant only in PERK^-/-^ cells; blue, significant only in WT cells; yellow, significant in both cell types. Statistical significance: p□<□0.05.

We next examined proteins with increased light intensities, *i.e*., decreased clearance. For proteins in the orange group, because decreased clearance is expected to increase heavy peptide intensity, the observed decreased heavy intensities suggested decreased synthesis. After adjusting for the effect of cell division, only 63 out of 710 proteins remained above the adjusted threshold of 0.95, indicating that reduced cell proliferation, rather than decreased protein degradation, accounts for most of the apparent slowing in protein clearance (**Fig. 4B**). Decreased protein synthesis and decreased clearance resulted in no significant change in total protein abundance. However, changes in protein synthesis could not be determined for proteins in the dark pink group, because reduced clearance would also contribute to increased heavy intensities. Notably, proteins in dark pink groups diverged into two distinct groups based on their H/L ratio changes (**Fig. 4A**). Because the H/L ratio reflects the relative proportion between newly synthesized and pre-existing protein pools, this subdivision might suggest that proteins within this group differ in the relative contributions of synthesis and clearance changes to their total abundance increase. To test it, we performed GO analysis and found that proteins with increased H/L ratios under Th treatment were enriched in amino acid metabolic and biosynthetic processes **(Fig. 4F),** consistent with previous evidence that they are transcriptionally regulated by ATF4^41,42^. After adjusting for the effect of cell division, the number of proteins enriched in these pathways decreased markedly **(Fig. S6C)**. Several proteins involved in the response to ER stress, including THBS1/HSPA5/CTH/NIBAN1/HYOU1, exhibited reduced degradation (**Fig. S6C, 4B**), indicating a synergetic regulation through both increased protein synthesis and reduced degradation. These findings suggest H/L changes provide additional information on determining protein clearance and synthesis regulation. Similarly, proteins in yellow group exhibited decreased light intensity, indicating increased clearance, which attributes to increased degradation (**Fig. 4B**); however, synthesis changes could not be determined because enhanced clearance alone would also reduce heavy intensity in the absence of additional regulation. Altogether, these results showed that protein synthesis and clearance can act independently or synergistically to modulate protein abundances, and different protein categories may rely on distinct regulatory mechanisms, with either synthesis or clearance serving as the driver of protein abundance changes.

It has been demonstrated that PERK mediates systematic translation inhibition and cell cycle arrest, which will lead to decreased protein synthesis and decreased clearance during UPR, respectively^43,44^. To examine whether the decreased synthesis and clearance observed in the orange group are regulated by PERK, we applied ISDia to PERK^-/-^ cells (**Table S4**) and examined how ER stress modulates protein synthesis and clearance in the absence of PERK. The knockout of PERK was validated by the absence of PERK protein expression in Western blotting analysis (**Fig. 4H**). In contrast to WT cells, ATF4 levels did not increase in PERK^⁻/⁻^ cells during the UPR, indicating disruption of the PERK-eIF2α-ATF4 axis. However, XBP1 and BiP are downstream targets of the other two ER stress senor in the UPR, IRE1α and ATF6, respectively. They were still upregulated under UPR conditions (**Fig. 4I**), confirming activation of the remaining arms of the UPR. ISDia quantified light, heavy peptide intensities, protein abundance and H/L ratios, for over 9,000 proteins across each treatment condition in PERK^-/-^ cells (**Fig. S7A**). Compared to WT cells, Tm and Th treated PERK^-/-^ cells showed fewer changes in heavy, light peptide intensities and H/L ratios (**Fig. 2F, S7B**). Changes under Th and Tm treatment were highly correlated (**Fig. S7C**). The Th versus DMSO comparison in PERK^-/-^ cells was highlighted in the subsequent analyses. Then we performed the pairwise comparison of changes in heavy, light, H+L and H/L ratio between Th and DMSO treatments **(Fig. 4J, S7D-E)**. Specifically, there are fewer proteins that have decreased clearance and synthesis under Th treatment in PERK^-/-^ cells than in WT cells (Group C in **Fig.4C**, **Fig. S7E**). These proteins showed less changes in clearance and synthesis in PERK^-/-^ cells **(Fig. 4K)**, indicating that PERK mediates both the suppression of protein synthesis and the reduced protein clearance associated with slower cell division. We performed GO enrichment and found that these proteins were enriched in RNA splicing, localization and transport (**Fig. S7F**).

### Differential regulation of protein synthesis and clearance across cellular compartments during the UPR

Organelles and large protein complexes are often equipped with distinct quality control and proteolytic mechanisms that maintain their abundances and functions^45–49^, leading us to investigate the regulatory mechanisms of protein abundance within these cellular compartments (CC). We found that different CCs exhibited diverse changes in heavy, light and H+L intensities, as well as H/L ratios under Th treatment in WT cells (**Fig. 5A**). Among all the CCs, chromatin proteins had the greatest degree of decreased heavy intensity. Mitochondrial proteins showed relatively less decreased heavy intensities compared to other CCs in WT cells, suggesting that the protein synthesis was less inhibited compared to other CCs. In addition, we observed that PERK^-/-^ cells showed smaller differences across compartments in changes on heavy, light, and H+L intensities, as well as in the H/L ratio compared with WT cells, indicating a less compartment-specific response (**Fig. 5B**). To further explore the compartment-specific regulation, we then compared changes in the light and heavy intensity under Th treatment relative to DMSO between WT and PERK^-/-^ cells (**Fig. 5C, Fig. S8A**). Across all compartments, the increase in light peptide intensities was significantly reduced in PERK^-/-^ cells **(Fig.S8A)**. Interestingly, only mitochondrial proteins showed a significantly smaller decrease in heavy peptide intensity changes in WT cells than in PERK^-/-^ cells. (**Fig. 5C)**, indicating that the regulation of mitochondrial protein synthesis during the UPR is PERK-dependent. A recent study has shown that PERK-ATAD3A interaction created a localized environment that allows translation of nuclear-encoded mitochondria proteins^50,51^. To assess whether that is the regulatory mechanism, we compared the protein level of ATAD3A in WT and PERK^-/-^ cells. ATAD3A levels increased in WT cells but not in PERK^-/-^ cells during the UPR (**Fig. 5D**), implicating its involvement in protecting mitochondrial protein synthesis in UPR is regulated by PERK.

**Figure 5.**
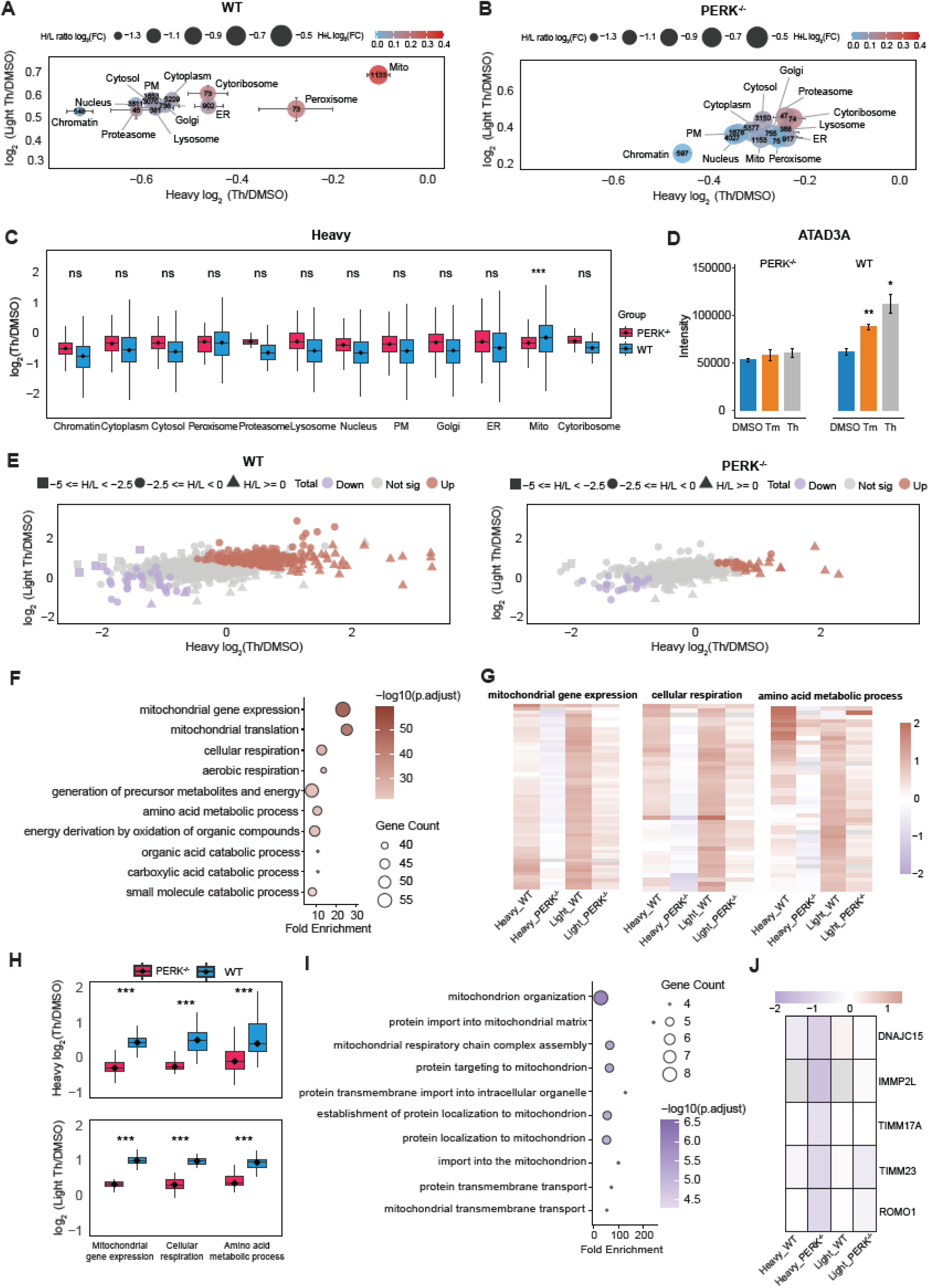
Differential regulation of protein synthesis and clearance across cellular compartments in WT and PERK^-/-^ cells during the UPR. A. Bubble plots showing the median of log_2_(fold change) in light, heavy, H+L, and H/L ratio across subcellular compartments in WT cells under Th treatment relative to DMSO. Numbers indicate the number of quantified proteins in each compartment. Error bars represent the standard deviation. B. Same plot as in (A), but for PERK^-/-^cells. C. Boxplot of log_₂_(fold change) in heavy peptide intensities across subcellular compartments in WT and PERK^-/-^ cells. P values are from paired, one-tailed t-tests. ***□p□<□0.001; **□0.001□<□p□<□0.01; *□0.01□<□p□<□0.05; ns, p□>□0.05. D. Changes of ATAD3A H+L intensity in WT and PERK^-/-^ cells under DMSO, Th and Tm treatments. ***□p□<□0.001; **□0.001□<□p□<□0.01; *□0.01□<□p□<□0.05. E. Bubble plots showing log_₂_(fold change) in light, heavy, H+L, and H/L ratios, for mitochondrial proteins in WT and PERK^-/-^ cells following Th treatment relative to DMSO. F. Biological processes enriched among mitochondrial proteins with significantly increased total abundance in WT cells following Th treatment relative to DMSO (H+L log_2_(Th/DMSO) > 0.58 & p-value < 0.05). G. Heatmap of heavy and light intensity changes for proteins belonging to selected enriched mitochondrial categories from **F**, including mitochondrial gene expression (GO:0140053), cellular respiration (GO:0045333), and amino acid metabolic process (GO:0006520). H. Boxplot of log_₂_(Th/DMSO) fold changes in heavy and light peptide intensities comparing WT and PERK-/- cells across categories in **G**. *P* values were calculated using paired, one-tailed t-tests. ***□p□<□0.001; **□0.001□<□p□<□0.01; *□0.01□<□p□<□0.05; ns, not significant. I. Biological processes enriched among the mitochondrial proteins with significantly decreased total abundance in PERK^-/-^ cells (H+L log_2_(Th/DMSO) < -0.58 & p-value < 0.05). J. Heatmap of heavy and light intensity changes for proteins annotated to the enriched protein targeting to mitochondrion (GO:0006626) category from Fig. 5I in WT and PERK^-/-^ cells.

Furthermore, we observed that more mitochondrial proteins exhibited significant total abundance changes in WT cells than in PERK^-/-^ cells (**Fig. 5E**). GO analysis of proteins with increased total abundance enriched for different categories in these two cell lines (**Fig. 5F, Fig. S8B**). To further examine how proteins with increased abundance in WT cells were impacted by PERK, we selected three GO categories, mitochondrial gene expression (GO:0140053), cellular respiration (GO:0045333) and amino acid metabolic process (GO:0006520), and compared the heavy and light intensity changes between the two cell lines. Proteins in each GO category showed increased light intensities in both WT and PERK^-/-^ cells, with significantly greater increases in WT cells, indicating stronger reduction of protein clearance **(Fig. 5G-H)**. Their corresponding heavy intensities were reduced in PERK^-/-^ cells under Th treatment, suggesting decreased protein synthesis during the UPR, whereas the heavy intensities in WT cells increased. We then examined the proteins with decreased total abundances and found proteins with decreased total intensities in WT were enriched for mitochondrial organization pathways (**Fig. S8C**). The result suggested elevated mitochondrial stress, as well as impaired mitochondrial organization and maintenance pathways, including fission-related processes^50,52–54^. In addition, we identified a group of proteins exhibiting decreased total intensities in PERK^-/-^ cells but not in WT cells. These proteins were significantly enriched for mitochondrial matrix protein import-related categories (**Fig. 5I**) and exhibited significantly reduced protein synthesis only in PERK^-/-^ cells but not in WT cells, suggesting that PERK is required for maintaining mitochondrial protein import machinery under stress condition **(Fig. 5J)**. Altogether, our results provide quantitative evidence that PERK regulates mitochondrial hemostasis during the UPR through tight control of protein synthesis and degradation.

We next focused on ER proteins. Unlike mitochondria proteins, ER proteins displayed largely similar changes in WT and PERK^-/-^ cells **(Fig.S9A)**. GO analysis indicated that proteins with significant increased total abundance in both cell lines are enriched in biological processes related to ER stress, ERAD pathway and protein folding (**Fig. S9B-C**). We observed that these proteins exhibited similar heavy and light intensity regulatory patterns in WT and PERK^-/-^ cells **(Fig.S9D)**, providing further proteomics evidence that these major pathways in ER proteostasis are regulated mainly through PERK-independent mechanisms, including the IRE1α and ATF6 pathways^3,55–58^. Interestingly, sterol metabolism-related proteins were significantly downregulated in WT but not PERK^-/-^ cells **(Fig. S9E)**, pointing to a potential PERK-dependent mechanism for future investigation.

### Regulation of synthesis and clearance across protein complex subunits during the UPR

Proteins often function as components of protein complexes that are co-regulated in response to environmental or intracellular changes. The composition and abundances of these complexes can impact their overall complex functions, and the state of a cell can be affected by the abundance or activity of protein complexes^21,59^. While previous studies have demonstrated that interacting proteins involved in protein complexes can be co-regulated^8,18,60^, how they are modulated through protein synthesis and clearance remains largely unexplored. To address this question, we used the protein complexes annotated in Ori *et al*^61^ and investigate their changes in protein synthesis and clearance during the UPR (**Table S5**). The majority of protein complexes exhibited decreased synthesis and clearance under UPR-inducing conditions in WT cells. However, a subset of complexes, including the presequence translocase-associated motor (PAM) complex, p97/VCP-VIMP-DERL1-DERL2-HRD1-SEL1L complex and ubiquitin ligase ERAD-L complex, displayed a synergistic increase in heavy and light peptide intensity under Th treatment (**Fig. 6A**).

**Figure 6.**
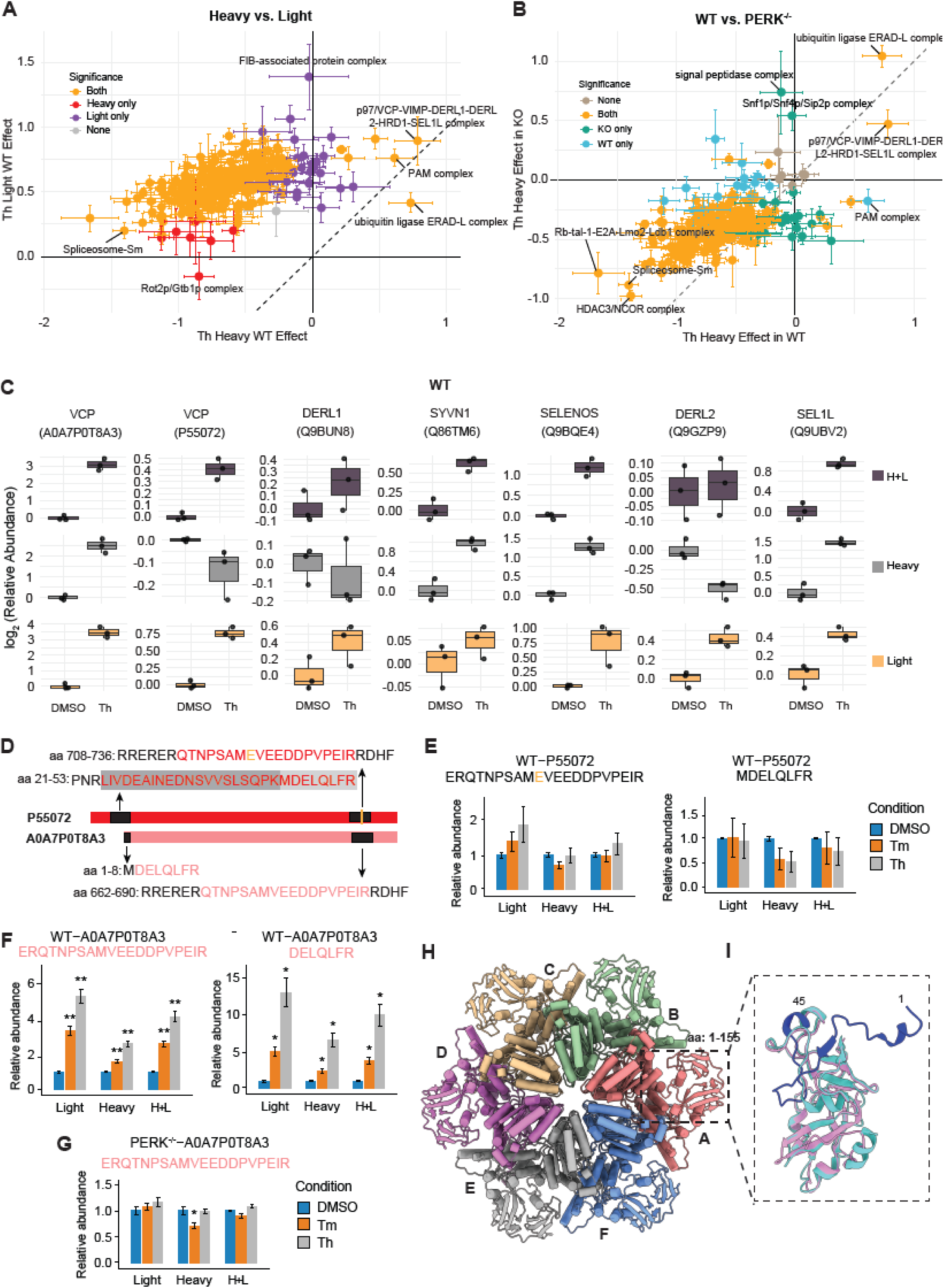
Differential regulation of synthesis and clearance among subunits across and within protein complexes during the UPR. A. Scatter plot showing the changes in heavy and light intensities for protein complexes in WT cells treated with Th relative to DMSO. Statistical significance was defined as an adjusted p-value < 0.05. Error bars represent the standard error. B. Scatter plot showing the changes in heavy peptide intensities for protein complexes under Th treatment relative to DMSO in WT and PERK^-/-^ cells. Statistical significance was defined as an adjusted p-value < 0.05. Error bars represent the standard error. C. Boxplots showing changes in heavy, light and H+L intensities for subunits of p97/VCP-VIMP-DERL1-DERL2-HRD1-SEL1L complex in WT cells under Th treatment relative to DMSO. D. Schematic showing isoform-specific peptides for the two p97/VCP isoforms. E. Relative abundance of long p97/VCP isoform-specific peptides (UniProt ID: P55072) under DMSO, Th, and Tm treatment in WT cells. K. Same plots as in E for short p97/VCP isoform-specific (UniProt ID: A0AP0T8A3) peptides in WT cells. ***□p□<□0.001; **□0.001□<□p□<□0.01; *□0.01□<□p□<□0.05. L. Relative abundance of short p97/VCP isoform-specific peptides under DMSO, Th, and Tm treatment in PERK^-/-^ cells. ***□p□<□0.001; **□0.001□<□p□<□0.01; *□0.01□<□p□<□0.05. F. AlphaFold3-predicted hexameric model of the short p97/VCP isoform (UniProt ID: A0AP0T8A3), shown in top-down view with each protomer colored distinctly (labeled A-F). The region corresponding to residues 1-155 of the short isoform is indicated. G. Structural alignment of the N-terminal domains of the short isoform (pink, residues 1-155) and long isoform (cyan, residues 1-200), highlighting the additional N-terminal extension (residues 1-45, dark blue) present only in the long isoform. This extension includes the first β-strand that contributes to a β-sheet, potentially stabilizing the overall N-terminal fold.

The PAM complex is a mitochondrion complex essential for importing preproteins into the mitochondrial matrix^62–64^. The Th-induced increase in heavy intensities for PAM complex subunits observed in WT cells diminished, whereas the light intensities reduced in PERK^-/-^ cells, indicating that its regulation is PERK-dependent (**Fig. 6B, Fig. S10A**). In contrast, the upregulation of heavy and light intensities for both the p97/VCP-VIMP-DERL1-DERL2-HRD1-SEL1L complex and the ubiquitin ligase ERAD-L complex were also observed in PERK^-/-^ cells (**Fig. 6B, Fig. S10A**), suggesting PERK-independent regulation. These two complexes play key roles in ERAD^4,65,66^, which can be further activated by the ATF6 and/or IRE1α-XBP1 during the UPR^3,55–58^. Interestingly, we also observed significant downregulation in the synthesis of spliceosome components, implying global alterations in RNA splicing during ER stress (**Fig. 6B**). Despite substantial changes in the sum of heavy and light intensities, changes in H/L ratios were minimal under Th treatment compared to DSMO in WT cells (**Fig. S10B**).

To determine whether all subunits within a complex are regulated uniformly, we analyzed the regulation of individual components of the p97/VCP-VIMP-DERL1-DERL2-HRD1-SEL1L complex in WT cells (**Fig. 6C**). This complex is known to be transcriptionally upregulated under stress conditions^36,67^. Integrated with the observed changes in heavy and light intensities, it provided insights into the underlying mechanisms. The RING finger E3 ubiquitin ligase SYVN1 (HRD1), its cofactor SEL1L, a scaffold protein that stabilizes HRD1 and facilitates substrate recognition^66^, and SELENOS (VIMP), which recruits p97/VCP to the ER membrane and coordinates its interaction with other ERAD components, all showed significantly increased abundance with increased heavy and light intensities. This evidence suggests that the increased abundance is driven by decreased clearance and enhanced synthesis. DERL1 and DERL2, which form the retrotranslocation channel for ERAD substrates^68,69^, remained largely unchanged. Strikingly, we found that the two isoforms of p97/VCP (A0A7P0T8A3 and P55072), a transitional ER ATPase that supplies energy for protein extraction during ERAD, were also regulated by different synthesis and clearance pattern during the UPR (**Fig. 6C**). All these results showed that the abundance of subunits within a complex could be regulated differently.

To more accurately investigate the regulation pattern for the two isoforms, we used isoform-specific peptide quantification (**Fig. 6D**) and found that peptides from the full-length isoform showed no significant changes in heavy and light intensity (**Fig. 6E**), whereas those from the shorter isoform exhibited significantly increased heavy and light intensity in WT cells (**Fig. 6F**). While the regulation of other subunits within the complex was conserved in PERK^-/-^ cells, the upregulation of the short VCP isoform was absent (**Fig. 6G, Fig. S10C**), indicating that its regulation is PERK-dependent. To validate the isoform-level change quantified by ISDia, we utilized parallel reaction monitoring (PRM)^70^ targeting isoform-specific peptides and quantified their fragment-ion intensities in DMSO and Th conditions in both WT and PERK^−/−^ cells. PRM recapitulated the ISDia trends across light, heavy, and total intensities, further supporting our conclusion that the short VCP isoform is upregulated under ER stress in WT but not in PERK^-/-^ cells (**Fig. S11 A-E**).

To investigate potential structural differences between the two isoforms of p97/VCP, we used AlphaFold3^71^ to generate a hexameric structural model of the short isoform (UniProt ID: A0AP0T8A3) (**Fig. 6H, Fig. S10D**). The predicted structural model revealed an overall organization highly similar to the previously reported cryo-EM structures of the long isoform^72,73^, consistent with conserved architecture in the AAA+ ATPase core domains. To examine isoform-specific features, we performed a structural alignment of their N-terminal regions (residues 1-200 in the long isoform and 1-155 in the short isoform). The alignment revealed that the long isoform contains an additional N-terminal segment (residues 1-45, shown in blue in **Fig. 6I**) that includes the first β-strand. This β-strand participates in an extended β-sheet packing, suggesting a potential role in stabilizing the N-terminal domain conformation of the long isoform (**Fig. 6I**). Given that the N-terminal region of p97/VCP is implicated in the binding of ubiquitin ligases and DUBs or other cofactors^74,75^ and involved in diseases conditions^76^, these structural differences raise the possibility that the two isoforms may exhibit differential binding specificity or regulatory interactions under various cellular stress conditions. Further biochemical and functional studies will be needed to determine whether the distinct N-terminal features contribute to isoform-specific roles in cellular proteostasis during the UPR.

In summary, our results provided preliminary support that protein complexes were regulated through coordinated synthesis and clearance, and that individual subunits within a complex can exhibit differential regulatory patterns. Using the ISDia method, we also uncovered PERK-dependent, isoform-specific regulation of VCP in modulating ERAD pathway during the UPR, providing new hypotheses for further mechanistic investigation.

## DISCUSSION

In this study, we established ISDia as a streamlined workflow that enables tracking and comparison of protein synthesis and clearance among multiple conditions. This approach offers several advantages over previous methodologies. First, rather than assuming that stressed cells remain in a steady state and deriving kinetic rate constants, we developed an alternative strategy that directly quantifies relative changes in protein clearance and synthesis by comparing light- and heavy-channel intensities across samples. It is particularly well-suited to capture proteome remodeling under non-steady-state conditions such as ER stress.^2,43,44^ Second, compared to previous methodologies that tracks protein synthesis and degradation, ISDia streamlined the workflow. Compared to mPDP^17^, it eliminates the need for multiplex labeling and fractionation. Compared with the time-resolved pSILAC-DIA workflow, ISDia reduces the number of samples analyzed and the required instrument time while maintaining high proteome coverage and robust quantitation, making it well suited for integration into high-throughput workflows^77,78^. Third, by integrating different timepoints, ISDia improves quantitation precision arising from peptides with extreme H/L ratios and insufficient signal in either channel (**Fig. 2D, 3C**), thereby improving quantitative accuracy and expanding proteome coverage. Finally, we demonstrated that cross-sample comparison of heavy and light intensities measured in DIA after normalization can recapitulate the theoretical difference of heavy and light intensity in standards on both Orbitrap Astral MS and Q-Exactive HF-X tandem MS. These results demonstrated that ISDia is widely applicable and not limited to instruments with high speed. Despite the proteome coverage might be lower **(Fig.S1C)**, it provides accurate cross-run quantitation. These features collectively highlight ISDia as a broadly applicable and scalable platform to track protein synthesis and degradation, respectively.

Besides these advantages, ISDia is designed to measure integrated changes in protein synthesis and clearance across the pooled labeling window under different conditions. Transient or biphasic responses may be attenuated or masked during pooling. Time-resolved analysis remains the preferred strategy when the timing and dynamics are of protein responses are of interest. Regarding the pooling scheme selection, we found pooling more time points increases proteome coverage, but the timing and number of pooled samples should be selected according to the expected dynamics, labeling duration, and experimental objective. For example, an earlier pooling window is more appropriate for capturing early responses, whereas a later or broader window is better suited to detecting sustained or progressively accumulating changes. For studies focused on proteome-wide responses rather than individual proteins, we recommend pooling samples from both early and late time points to improve measurement robustness while maintaining broad proteome coverage. In addition, because pooled ISDia signal reflects both baseline protein abundance and the process of interest, comparisons between different cell lines can be confounded by pre-existing differences in protein abundance. ISDia is therefore most reliable in matched, within-cell-line comparisons (e.g., treatment vs. vehicle), where cells share the same starting population and baseline abundances are expected to be equivalent at the onset of labeling. A further simplified implementation of ISDia, *i.e*, single time point pSILAC-DIA, would be used to measure light and heavy intensities at a single labeling time point, which could still provide insight into the relative contributions of protein synthesis and clearance to changes in protein abundance across conditions.

Using ISDia, we uncovered diverse regulations between protein synthesis and clearance during ER stress. By integrating heavy, light and total intensities, we examined protein synthesis and clearance and found that they can act either independently or synergistically to shape proteome remodeling during UPR. Distinct functional classes of proteins relying on different regulatory mechanisms to determine their abundance during the UPR. Notably, similar changes in total protein abundance can arise from distinct combinations of protein synthesis and clearance. These findings highlight the value of ISDia in resolving regulatory layers that are not apparent from total abundance measurements alone and establish a quantitative framework for future mechanistic studies. Although we aimed to establish a streamlined workflow for measuring protein synthesis and clearance without requiring cell division tracking, cell proliferation can be monitored to derive a correction factor that distinguishes changes in protein degradation from dilution effects mediated by cell division (**Fig. 4B**, **Method**). In this study, dividing cells were labeled for 48h under different conditions here. Under this experiment design, heavy peptide intensities represent the accumulative effects of both protein synthesis and clearance, while protein clearance itself is influenced by both degradation and cell division. Therefore, synthesis regulation could not be determined for certain groups (dark pink, yellow). Future application of ISDia in non-dividing cells or over shorter labeling periods may reduce the contribution of cell division mediated dilution and improve the resolution of synthesis and degradation changes across conditions.

We then used ISDia to investigate how PERK impacts protein synthesis and clearance during the UPR. Our results demonstrated that proteins exhibiting reduced synthesis and clearance in WT cells showed minimal changes in PERK^-/-^ cells (**Fig. 4K**), suggesting that the global protein synthesis and clearance decrease is PERK-dependent during UPR. This is consistent with the established role of PERK in driving global translational suppression and cell cycle arrest in response to ER stress^43,44^. In addition, our analyses of cellular compartments support a model that PERK drives a remodeling of the mitochondrial proteome during the UPR. In WT cells, PERK signaling may promote retention of pre-existing mitochondrial proteins by decreasing protein clearance^79^ while simultaneously protecting mitochondrial protein translation from the global translation inhibition through the PERK-ATAD3 interaction^51^. In the absence of PERK, other mechanisms, including compensatory eIF2α phosphorylation and chronic suppression of mTORC1 signaling, may regulate protein synthesis and clearance, resulting in distinct regulatory patterns^2,5,44,80,81^. In mitochondria, PAM complex engages TIM23 complex to couple translocation across the inner membrane and import proteins into the mitochondria matrix^82–84^. We found that PAM complex was upregulated in WT cells but not in PERK^-/-^ cells, while the TIM23 complex (**Fig. S10B**), including TIMM23, ROMO1 and TIMM17A^85,86^ (**Fig. 5J**), was maintained in WT cells but markedly reduced in PERK^-/-^ cells. These findings suggest PERK maintained mitochondrial protein import during ER stress. In contrast, ER proteins show comparatively minimal PERK-dependent changes, which is consistent with the literatures that ER proteostasis is mainly regulated by IRE1α and ATF6 during the UPR^3,55–58^. Furthermore, we identified isoform-specific regulation of VCP mediated by PERK in response to UPR activation. This compelling example demonstrated that ISDia offers a powerful platform to investigate such isoform-specific regulation with high proteome coverage. Altogether, these results provide an unprecedented, system level view of global protein homeostasis control, highlighting the diverse strategies that cells employ to adapt to stress conditions and facilitate follow-up mechanistic investigation. These findings were independently validated using both Th (**Fig. 4-6**) and Tm treatments (**Fig. S12-14**). Their reproducibility across two mechanistically distinct ER stressors indicates that they reflect a core PERK-dependent adaptive program rather than compound-specific effects, thereby strengthening the generalizability of the biological conclusions.

In summary, we developed ISDia, a streamlined proteomics platform, leveraging direct light and heavy intensity comparison across samples in pSILAC-DIA to elucidate the regulation of protein synthesis and clearance. We uncovered diverse regulatory mechanisms governing protein abundances during the UPR, driven by distinct combination of protein synthesis and clearance. These results highlight the potential of ISDia as a powerful tool for providing insights into protein abundance regulatory mechanisms in many contexts, including drug development and neurodegenerative disease models.

### Limitations of the study

It is known that ER stress induces protein aggregation^3,87^, and it is possible that the newly synthesized proteins are misfolded during the UPR and will undergo more rapid clearance. Coupling ISDia with differential buffers for protein extraction or strategies to barcode proteins based on their time of synthesis^88^, could enable the separation of misfolded proteins to show how synthesis and clearance contributes to their accumulation, particularly under different genetic background or disease models.

Despite ISDia enabling protein abundance regulation analysis under non-steady states, it cannot be directly applied to clinical samples, as it requires the incorporation of isotope-labeled amino acids, as is possible in model systems, such as mouse and fly.

Elucidating mechanisms of protein synthesis and degradation regulation, such as identification of upstream E3 ligases, or defining the roles of proteins involved in the UPR, is beyond the scope of this study. To fully understand this mechanism, further studies are needed.

## METHODS

### Cell Culture and Isotope Labeling

Both Hela and RPE1 cells were cultured in DMEM supplemented with 10% fetal bovine serum (FBS) and 1% penicillin-streptomycin (P/S) at 37°C in a humidified incubator with 5% CO_₂_. For heavy isotope labeling, cells were gradually adapted to DMEM supplemented with 10% dialyzed FBS and 1% Penicillin-Streptomycin (P/S). For MS standards with pre-defined heavy/light (H/L) ratios, HCT116 cells were passaged seven times in SILAC DMEM supplemented with heavy isotope-labeled arginine (+10.008269 Da) and lysine (+8.014199 Da), 10% dialyzed FBS, and 1% P/S. Corresponding light control cells were cultured in SILAC DMEM with native arginine and lysine under identical conditions. Fully heavy isotope-labeled cells and light control cells were harvested separately and stored at -80°C for subsequent proteomics analysis.

HeLa cells were treated with 2.5 µg/mL tunicamycin (Tm) and 200 nM thapsigargin (Th) for 0, 6, 12, 24, and 48 hours. Isotope labeling was initiated simultaneously using SILAC DMEM supplemented with heavy isotope-labeled arginine (+10.008269 Da) and lysine (+8.014199 Da), 10% dialyzed FBS, and 1% penicillin-streptomycin. DMSO-treated cells served as controls. Each condition was performed in triplicate. Following treatment, cells were harvested and stored at -80°C for proteomics analysis.

### Cell Confluency Measurements

Cells were seeded at a density of 3,000 cells/well in 96-well plates and allowed to adhere for 15 h prior to treatment (4 replicate wells per treatment group). Imaging was initiated immediately after treatment. Cell proliferation was monitored using an Incucyte SX5 live-cell imaging system (Sartorius) housed in a humidified incubator at 37°C and 5% CO_₂_. Phase-contrast images (10× objective, 1 field per well) were acquired every 3 h over a 4-day period. Whole-well images were analyzed using Incucyte software (version 2025B), and confluency was quantified using the Classic Confluence segmentation algorithm, with consistent parameters applied across all experimental groups within each experiment. Confluency values were averaged across replicate wells and used as a proxy for relative cell growth.

### Western Blotting

Cells were harvested and lysed using RIPA buffer (25 mM Tris-HCl pH 7.6, 150 mM NaCl, 1% NP-40, 1% sodium deoxycholate, 0.1% SDS) supplemented with 1x protease inhibitor cocktail. Lysates were centrifuged at 14,000 rpm for 10 minutes at 4°C, and the supernatant was collected for protein quantification using the BCA assay. A total of 20 µg of protein per sample was used. Electrophoresis was performed at 200 V for 35 minutes using NuPAGE™ Bis-Tris gels. Proteins were transferred to PVDF membranes using the Invitrogen iBlot2 Gel Transfer Stacks (IB24001, 0.2 µm pore size). Membranes were then blocked for 1 hour at room temperature in 5% non-fat milk prepared in TBST buffer (20 mM Tris-HCl pH 8.0, 137 mM NaCl, 0.05% Tween-20). Membranes were incubated overnight at 4°C with the following primary antibodies: monoclonal rabbit anti-PERK (1:1000), anti-BiP (1:1000), anti-ATF4 (1:1000), anti-XBP1s (1:1000), and mouse anti-β-Actin (1:2500). Membranes were washed with TBST and incubated for 1 hour at room temperature with appropriate HRP-conjugated secondary antibodies: anti-rabbit IgG for rabbit primary antibodies or anti-mouse IgG for β-Actin. Protein bands were visualized using the Pierce™ Enhanced Chemiluminescence (ECL) detection system (Thermo Fisher Scientific) on a BioRad imager.

### Sample preparation for proteomic analysis

Cell pellets were lysed in 8M urea, 200 mM EPPS, protease and phosphatase inhibitor (ThermoFisher Scientific) and sonicated. Then the supernatants were collected by centrifugation, and the protein concentrations were measured using Pierce BCA assay kits. Subsequently, proteins were reduced with 5□mM TCEP for 15 minutes, alkylated with 15□mM iodoacetamide in the dark for 30 minutes, and quenched with 10□mM DTT at room temperature. Next, 20 µg of protein was digested with LysC and trypsin overnight at 37°C with shaking at 1400 rpm on a ThermoMixer. Heavy-only or heavy-light mixed samples were prepared at predefined H/L ratios of 1:27, 1:9, 1:3, 1:1, 3:1, 9:1, and 27:1 prior to protein digestion, with three replicates per ratio. For ISDia, samples from the same treatment across four time points were pooled before digestion, also with three replicates per group. The samples were then desalted, dried and resuspended in 5% acetonitrile (ACN) and 5% formic acid (FA) for mass spectrometry analysis.

### Liquid chromatography (LC)

LC was performed using a Vanquish Neo ultra high-performance liquid chromatography (UHPLC) system, configured in trap-and-elute format. Peptides were separated on an Aurora Frontier^TM^ TS C18 UHPLC column (IonOpticks, 60 cm length, 75 µm inner diameter, 1.7 µm particle size). During LC separations, mobile phase A (MPA) was 0.1% formic acid (FA) in water and mobile phase B (MPB) was 80% or 99.9% ACN in water with 0.1% FA. A 50-min active gradient ramped, at a flow rate of 0.3 μl/min, from 9.6% to 14.4% ACN from 0.1 to 8 min, 14.4% to 36% ACN from 8 to 50 min at a flow rate of 0.2 μl/min, 36% to 80% ACN from 50 to 50.1min, and held at 80% ACN to wash the column and return to 1% ACN for re-equilibration.

### Mass spectrometry data acquisition

For the DIA experiments on the Orbitrap Astral mass spectrometer, MS1 spectra were acquired at a resolution of 240,000 over an m/z range of 380–980. The MS1 normalized AGC target was set to 250%, and the maximum injection time was set to 50 ms. DIA MS2 scans were recorded at a resolution of 80,000 with a normalized AGC target of 500%. The HCD normalized collision energy was set to 25%, and a 0.6 s cycle time was used with a maximum injection time of 3.5 ms and a 2 Th isolation window. Equal amounts of sample were injected for all runs.

In DDA mode, MS1 spectra were acquired in the Orbitrap at a resolution of 180,000, with a normalized AGC target of 100% and a maximum injection time of 50 ms over an m/z range of 350–1350. The precursor fit window was set to 1.2 m/z, and dynamic exclusion was enabled with a 15 s exclusion duration after a single selection. The data-dependent method was set to select up to 30 precursors per cycle. MS2 spectra were acquired in the Astral analyzer with HCD fragmentation at a normalized collision energy of 25%, a normalized AGC target of 200%, and a maximum injection time of 15 ms. Equal amounts of sample were injected for all runs.

For PRM validation, we performed two acquisition experiments on the Orbitrap Astral mass spectrometer. Full-scan MS1 spectra were acquired in the Orbitrap analyzer at a resolution of 60,000 over an m/z range of 200–1500, using a normalized AGC target of 100% and a maximum injection time of 20 ms. Targeted MS2 (tMS2) scans were acquired in the Astral analyzer using a normalized AGC target of 1500% and a maximum injection time of 150 ms. The normalized HCD collision energy was set to 25%. Target precursor m/z values and charge states were specified in an inclusion table (**Table S6**). Equal amounts of sample were injected for all runs.

### Data analysis

#### DIA data analysis

DIA raw files were processed in Spectronaut (v19.6.250122.62635) using the default directDIA workflow and a human reference proteome database (UniProt release 2024-12-19; 83,401 entries). The same search settings were applied across all datasets and MS platforms. In this search settings, retention-time calibration was performed directly in Spectronaut without external iRT spike-in standards. We used the default high-precision iRT-to-RT calibration based on endogenous peptides, with deep-learning–assisted iRT calibration used as a fallback when empirical anchor information was insufficient.

For Pulsar search settings, Trypsin/P was selected as the digestion enzyme with up to two missed cleavages. SILAC labeling was defined as a two-channel setup, with no label assigned to Channel 1 and Arg10/Lys8 assigned to Channel 2. Carbamidomethyl (C) was specified as a fixed modification, while Acetyl (Protein N-term) and Oxidation (M) were included as variable modifications. In the Result Filters settings, the Top3–6 best fragments per peptide were used.

For DIA analysis settings, in the Identification panel, the precursor q-value and experiment-wide protein q-value thresholds were set to 0.01, and the run-wise protein q-value threshold was set to 0.05. In the Quantification panel, the MultiChannel Q-value filter was set to “Group Q-value”. Quantification was performed at the MS2 level, and cross-run normalization was enabled. The “Exclude All Multi-Channel Interferences” option was enabled, which excludes potentially interfering co-fragmented b-ions shared by heavy and light peptides. Peptide and protein quantities were summarized using a maximum of Top 3 precursors and Top 3 striped peptide sequences, respectively. The “Minimum Log2 Precursor Quantity” parameter was set to 0. In the Workflow settings, “MultiChannel Workflow Definition” was set to “From Library Annotation” and the “Fallback Option” was set to “Labeled.” PG. MS2 Channel 1 and PG. MS2 Channel 2 values are exported for each protein quantitation. Isoform specific peptides were used for A0AP0T8A3 quantitation.

#### DDA data analysis

For Heavy labeling efficiency analysis, DDA raw files were analyzed using FragPipe (version 22.1) with the default workflow settings. Briefly, a human reference database (UniProt 2024/12/19 release, containing 83,401 entries) with concatenated and decoys was applied. In the MSFragger, in addition to the common variable and fixed modifications, heavy isotope modifications on R (+10.008269□Da) and K (+8.014199□Da) were designated as variable modifications for heavy-labeled peptides. PTM site localization was carried out with PTMProphet with a minimum probability 0.5. Protein inference was performed with ProteinProphet and results were generated and filtered using a sequential strategy with PSM, peptide and protein-level FDR controlled at 1%. IonQuant was configured to quantify precursors based on the MS1 area, with intensities normalized across runs. MaxLFQ Intensity from “protein label quant” report were used for following data analysis.

#### PRM data analysis

It was performed in Skyline (version 25.1.0.237). Raw files were imported and peptides were configured in Peptide Settings with Carbamidomethyl (C) and Acetyl (Protein N-term) specified as structural modifications. Heavy isotope labeling was defined in Isotope Modifications as Lys (+8.014199 Da) and Arg (+10.008269 Da). In Transition Settings, targets were filtered to precursor charge states 2 and 3, and fragment ion types b and y. For full-scan extraction, MS1 filtering was set to include the first three isotope peaks, with the precursor mass analyzer set to centroided and mass accuracy set to 10 ppm. For MS/MS filtering, the acquisition method was set to PRM, with the product mass analyzer set to centroided and mass accuracy set to 10 ppm. Retention time filtering was configured to use all matching scans. Peak boundaries were inspected in Skyline and adjusted manually when necessary to ensure consistent integration across samples. Total area of fragments was used for quantitation.

#### Cellular compartment and protein complex analysis

Protein lists corresponding to different cellular compartments were obtained from UniProt based on Gene Ontology (GO) annotations (Downloaded in March 2025). The following GO terms were used to retrieve proteins assigned to specific cellular compartments: cytosolic ribosome (GO:0022626), nucleus (GO:0005634), cytosol (GO:0005829), mitochondrion (GO:0005739), golgi apparatus (GO:0005794), endoplasmic reticulum (GO:0005783), plasma membrane (GO:0005886), lysosome (GO:0005764), peroxisome (GO:0005777), proteasome complex (GO:0000502), chromatin (GO:0000785), and cytoplasm (GO:0005737). Only reviewed entries from homo sapiens were used for analysis. Annotations that mapped genes to protein complexes were obtained from Ori *et al.* ^61^, which compiled information from their own literature search, the CORUM protein complex database^89^, and the COMPLEAT data resource^90^. We mapped ENSEMBL gene IDs from the annotated protein complexes to gene symbols, which were then mapped to the protein data.

#### Detecting treatment differences in protein complex-wide intensity

We first filtered the protein data to protein complexes with three or more subunits observed in the data to avoid making inferences based on only one or two proteins. We used linear mixed effects models (LMMs) to test for a treatment effect on intensity shared across complex subunits. For each protein complex, we fit the following joint model (Equation 4) of the complex’s annotated proteins for a given cell background and treatment:

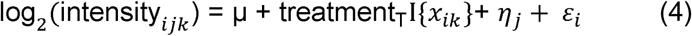

where intensity*_ijk_* is the intensity value for protein *j* of sample *i* exposed to treatment *k*, *μ* is a shared intercept, treatment_T_ is the effect of treatment Tm or Th, I{*x_ik_*} is an indicator function that sample *i* was exposed to treatment T, *η_j_* is a random effect that allows for models protein-specific deviations from the overall complex, and *ɛ_ijk_* is the unstructured random error term. The random terms are modeled as *η_j_* ∼ N(0, *τ*^2^) and *ɛ_i_* ∼ N(0, *σ*^2^) where *τ*^2^ and *σ*^2^ are variance components. The treatment effect was tested with ANOVA by comparing the model in Eq1 to the null model excluding it. The LMMs were fit using the R package lme4^91^. Because our focus was testing, maximum likelihood estimation was used instead of restricted maximum likelihood (REML). Statistical significance was determined based on false discovery rate (FDR)-adjusted p-values, using the Benjamini-Hochberg method^92^.

#### Statistical analysis

Protein-level light and heavy channel intensities were exported from Spectronaut, with MS2Channel1 assigned as the light channel and MS2Channel2 assigned as the heavy channel. For the calculation of total intensity (H+L) and H/L ratios, only replicates with both light and heavy intensity values available were included. H+L was calculated as the sum of light and heavy intensities, and the H/L ratio was calculated as heavy intensity divided by light intensity. Group means, coefficients of variation (CVs), and statistical tests were calculated only for proteins quantified in all three biological replicates under each condition. Unless otherwise specified, statistical significance between conditions was assessed using Welch’s two-sample *t*-test assuming unequal variance.

#### Estimating the cell-division offset for light-channel adjustment

To account for proliferation-driven differences in light-channel intensity, we derived a confluency-based offset and applied it prior to differential analysis.

For each condition *c* at time point *t*, relative growth was calculated as

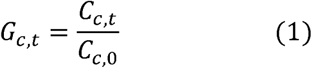

where *C_c,t_*, is the Incucyte-derived confluency for condition *c* at time *t*, and *C_c,_*_0_, is the confluency at the initial time point. The corresponding light-channel dilution factor was defined as the inverse of relative growth:

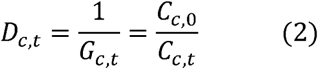

For pooled samples combining *K* time points (*t* ɛ *T*), a mean dilution factor was calculated as

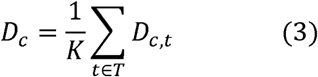

For each treatment condition *x*(Tunicamycin or Thapsigargin), an adjustment factor *M_x_* was defined relative to the DMSO control:

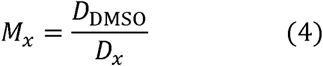

with *M*_DMSO_ = 1 by definition. This factor was applied to each biological replicate’s raw light-channel intensity (*L_x,r_*, *r* = 1, … , *n_x_*):

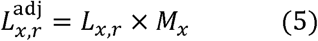

Adjusted replicate-level intensities *L_x,r_*^adj^ were averaged to obtain the mean adjusted intensity per condition,

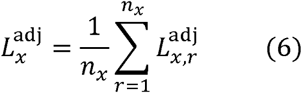

and adjusted log_2_ fold changes were calculated as

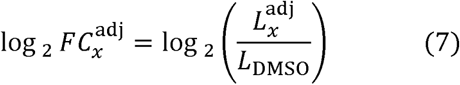

Because the same adjustment factor was applied uniformly to all replicates of a given treatment, the growth-adjusted significance thresholds shown on the raw (unadjusted) log_2_FC scale were shifted by the cell division offset *O_x_* = -log_2_(*M_x_*):

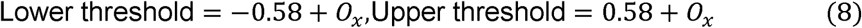

Adjusted light-channel *p* values were calculated using an unpaired, two-sided *t*-test comparing replicate-level adjusted treatment intensities (*L_x,r_*^adj^) against the corresponding DMSO intensities.

### Derivation of the pooled ISDia fold change

Let *T_i_* and *C_i_* denote the abundance of a given protein at time point *i* in the treatment and control conditions, respectively. The time-specific fold change *FC_i_* on the linear scale is defined as

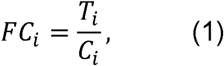

and therefore

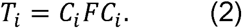

Because equal amounts of total protein from *n* time points are combined before MS analysis, the fold change measured from the pooled samples is

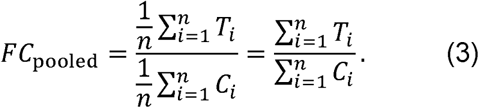

Substituting *T_i_* = *C_i_FC_i_* gives

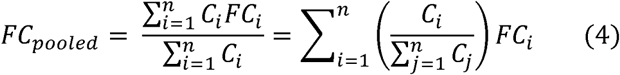

Defining the relative contribution of that time point to the pooled control signal

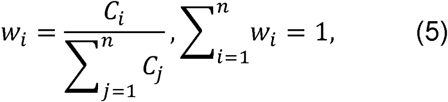

Equation (4) can be written as

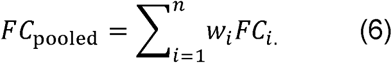

Thus, *FC*_pooled_ represents the weighted average of the time-specific fold changes, with each time point weighted according to its relative contribution to the pooled control signal.

## RESOURCE AVAILABILITY

### Lead contact

Requests for further information and resources should be directed to and will be fulfilled by the lead contact, Tian Zhang.

### Materials availability

This study did not generate new, unique reagents.

### Data and code availability

- The mass spectrometry proteomics data generated in this study have been deposited to the ProteomeXchange Consortium via the PRIDE^93^ partner repository with the dataset identifier PXD066472. Reviewer can access the dataset by logging in to the PRIDE website using the following account details: Username: Password: 2lv7idp1cTIh.
- All data processing and statistical analyses were conducted with the R statistical programming language^94^. All processed data and the R code used for statistical analyses and generating the reported findings are publicly available on GitHub (https://github.com/TianZhangLab/Code-for-Integral-Synthesis-and-clearance-analysis-via-DIA-ISDia). README file at the top of the directory provides description of directory contents, which includes an R script for generating each figure.
- Any additional information required to reanalyze the data reported in this paper is available from the lead contact upon request.

## DECLARATION OF GENERATIVE AI AND AI-ASSISTED TECHNOLOGIES IN THE WRITING PROCESS

During the preparation of this work, the authors used ChatGPT in order to improve language and readability. After using this tool, the authors reviewed and edited the content as needed and took full responsibility for the content of the publication.

## ACKNOWLDEGMENTS

We thank all members of the Zhang Lab for their active discussions and support. T.Z. was supported by NCI grant R00CA273170. J.A.P. was funded by NIH/NIGMS grants R01 GM132129 and R35 GM156406. We acknowledge funding support from the University of Virginia Comprehensive Cancer Center (UVACCC). We also thank Dr. Sina Ghaemmaghami from the Biology Department of University of Rochester, Dr. Luke Wiseman from the Department of Molecular and Cellular Biology at the Scripps Research, Drs. Chongzhi Zang, Yuh-Hwa Wang, Hui Li from the Department of Biochemistry and Molecular Genetics at the University of Virginia School of Medicine for their valuable suggestions and insightful discussion.

## AUTHOR CONTRIBUTIONS

T.Z. conceptualized the project. Y.D. and T.Z and designed the experiments. Y.D., D.K.Q, V. L., W. E. M., W. L., J.A.P. and T.Z. performed the experiments. J.Y. generated the structural model and performed structural analysis. Y.D., G.R.K. and T.Z. designed the statistical approach. Y.D., G.R.K. and T.Z. performed the analysis. Y.D., G.R.K., J.A.P., L.Q. and T.Z. wrote the manuscript. All authors reviewed the manuscript.

---

**Figure S1.**
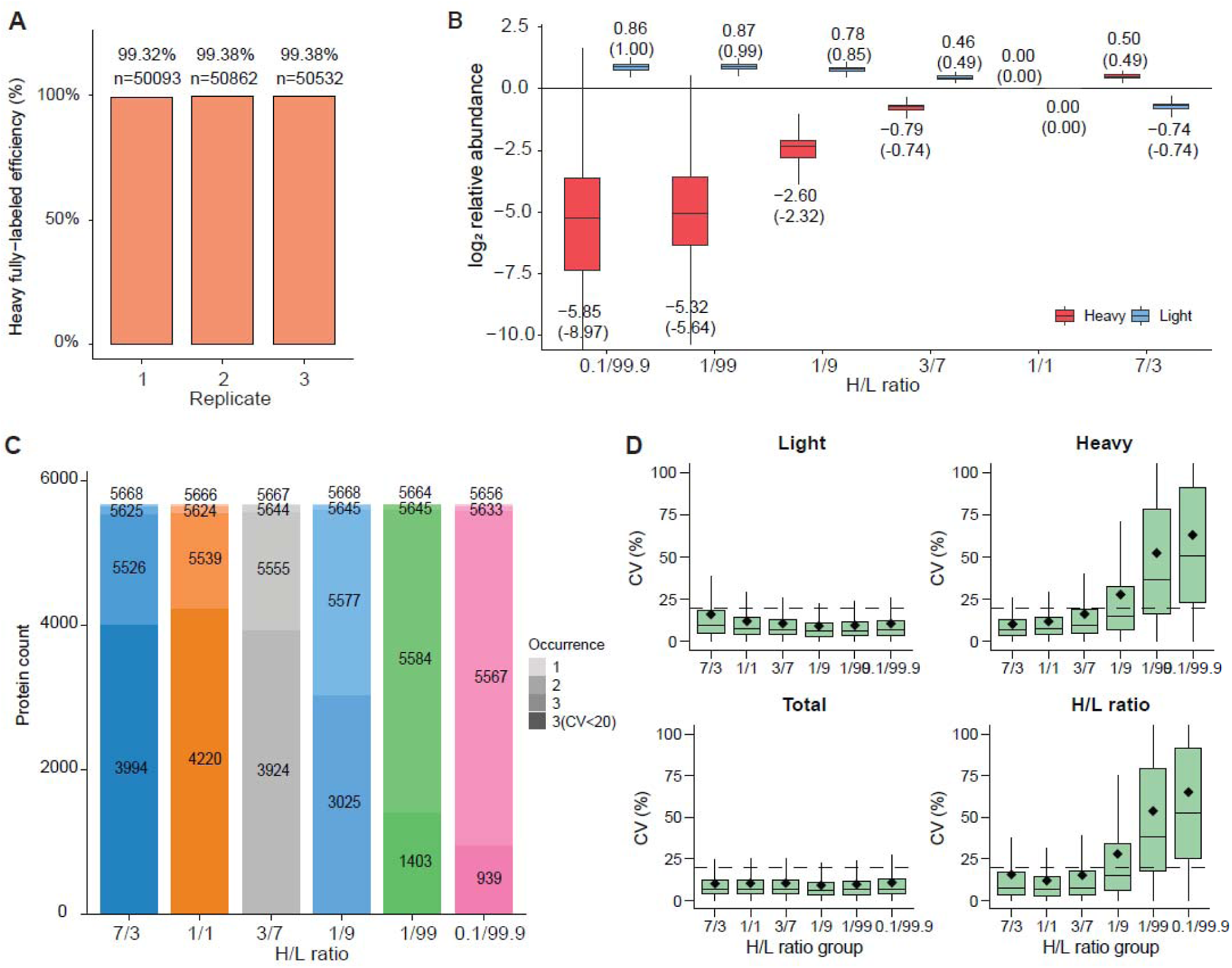
Assessment of labeling efficiency in HCT116 cells and quantitation of predefined H/L ratio standards using DIA on a Q Exactive HF-X MS, related to Figure 1. A. Heavy-labeling efficiency of HCT116 cells used to generate standards with pre-defined H/L ratios. Heavy-labeling efficiency was calculated as the intensity of the fully labeled peptide divided by the total peptide intensity (fully labeled + partially/unlabeled forms) for peptides containing lysine (K) or arginine (R). B. Cross-sample quantitation of light and heavy peptide intensities in standards acquired in DIA mode on a Q-Exactive HF-X tandem MS from Pino *et al*. Light and heavy intensities were normalized to the corresponding values from the 1:1 standard sample. Fold changes are labeled, with theoretical fold changes indicated in parentheses. C. Proteome coverage across standards shown in B. Each bar includes four numbers indicating the count of proteins for which both light and heavy intensities were detected in one, two, three replicates, or were consistently quantified across all three replicates with a coefficient of variation (CV) < 20% for light, heavy, total intensities, and H/L ratios. Only proteins were quantified with both heavy and light intensities in three replicates were included for CV calculation. Total intensity (H+L) and H/L ratio were computed when both heavy and light quantified in each replicate. D. Boxplots showing the CV of light, heavy, total peptide intensities and H/L ratios across three replicates of the standards. The black dashed line indicates the 20% CV threshold.

**Figure S2.**
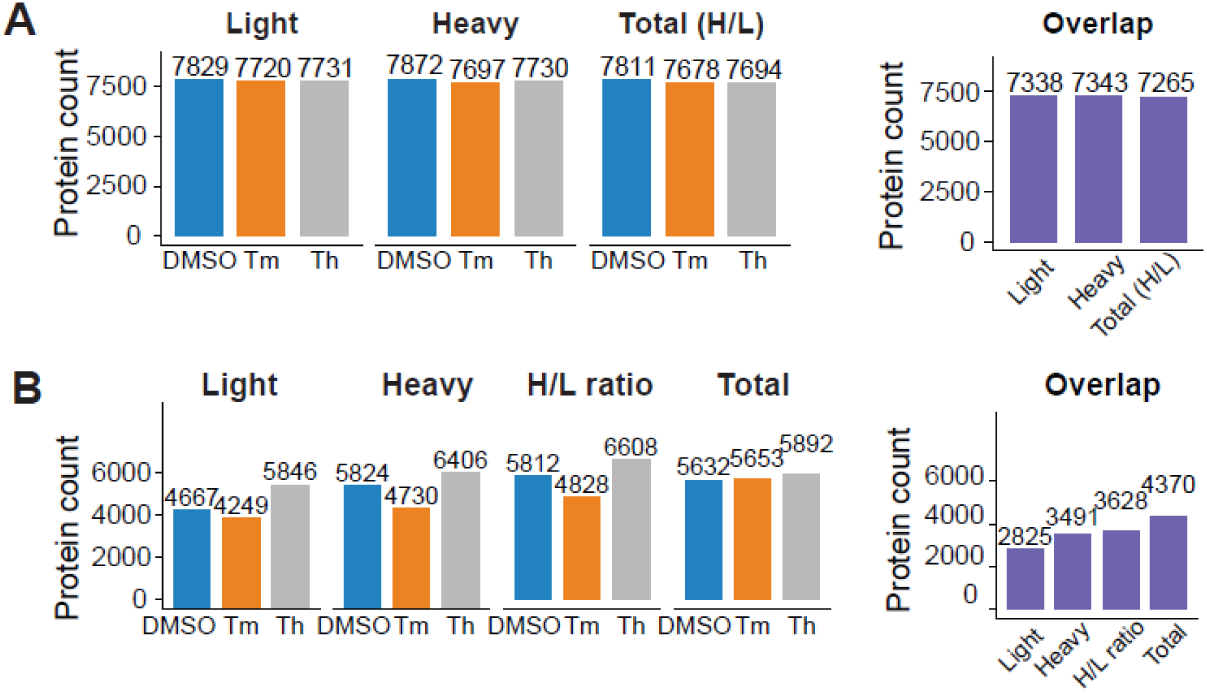
Proteome coverage and quantitation reproducibility of a single timepoint (48h) in pSILAC-DIA experiment. A. Number of proteins quantified in all three biological replicates at 48h under each condition. Bar plots show the number of proteins with quantified light, heavy, and total intensity (H/L ratio) in DMSO, Th, and Tm. The right panel shows the overlapping quantified proteins across three conditions for each metric. Total (H + L) and H/L ratio values are calculated only for proteins with both light and heavy intensities quantified in all three replicates. B. Same as A, after applying CV< 20% filtering.

**Figure S3.**
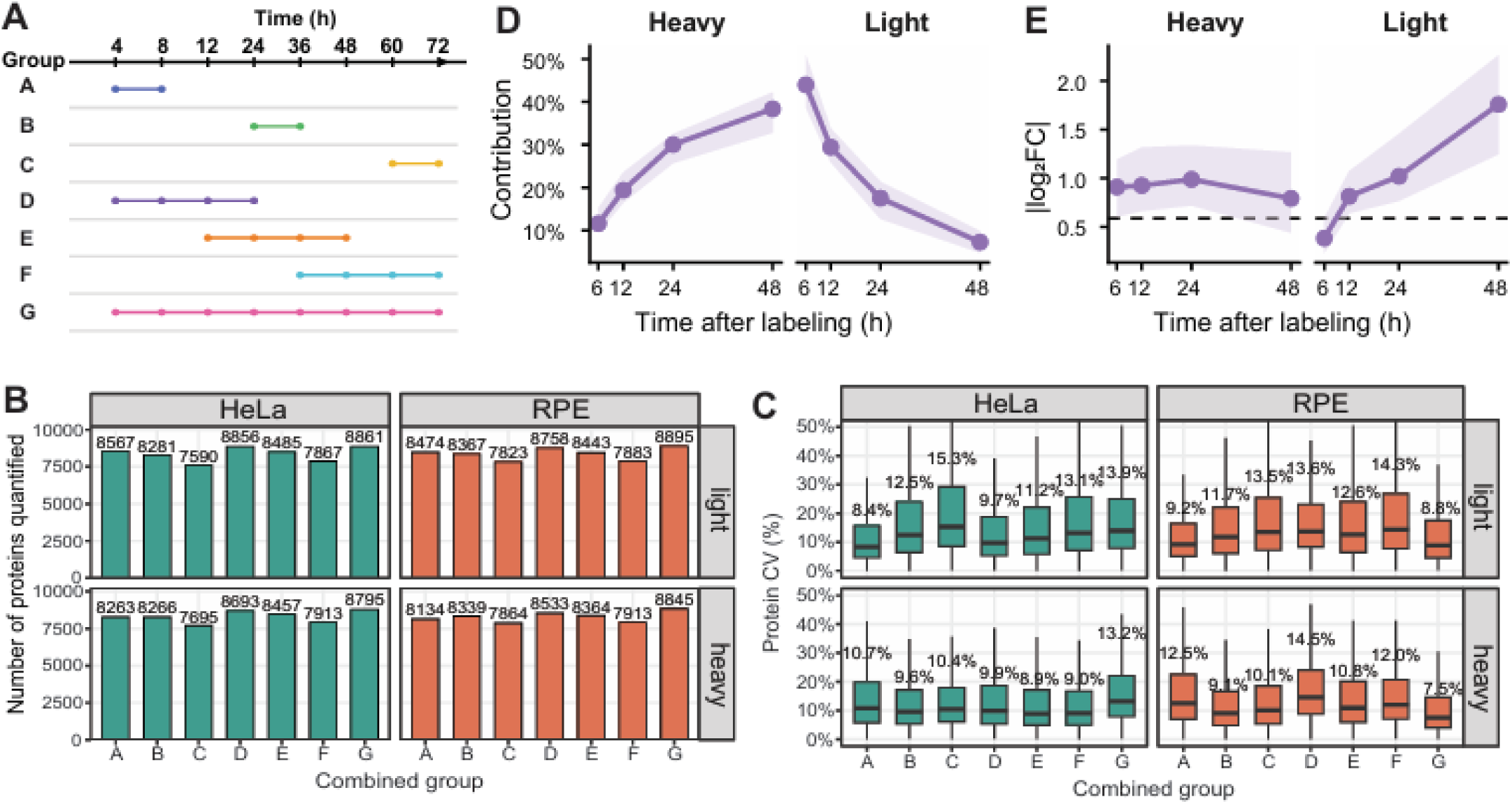
Method development of ISDia. A. Pooling schemes of HeLa and RPE1 pSILAC samples collected under steady-state conditions: A, 4 and 8 h; B, 24 and 36 h; C, 60 and 72 h; D, 4, 8, 12, and 24 h; E, 12, 24, 36, and 48 h; F, 36, 48, 60, and 72 h; and G, all eight time points. B. Numbers of light and heavy proteins quantified in all three biological replicates for each pooling scheme in HeLa and RPE1 cells. C. Distribution of protein-level CVs for light and heavy proteins quantified in all three biological replicates. Median CV values are indicated above each box. D. Relative contribution of individual time points to the pooled heavy and light intensity in DMSO samples. For each protein, the contribution of a time point was calculated as the mean intensity at that time point divided by the sum of the mean intensities across all four time points (6, 12, 24, and 48 h). Lines and points represent the median contribution across proteins, and shaded regions indicate the interquartile range. E. Temporal response trajectories of proteins in the heavy and light channels under Th treatment relative to DSMO control. Lines and points show the median absolute log2 fold change at each time point, and shaded regions indicate the interquartile range. The dashed horizontal line indicates the fold-change cutoff of |log_2_FC| = 0.58.

**Figure S4.**
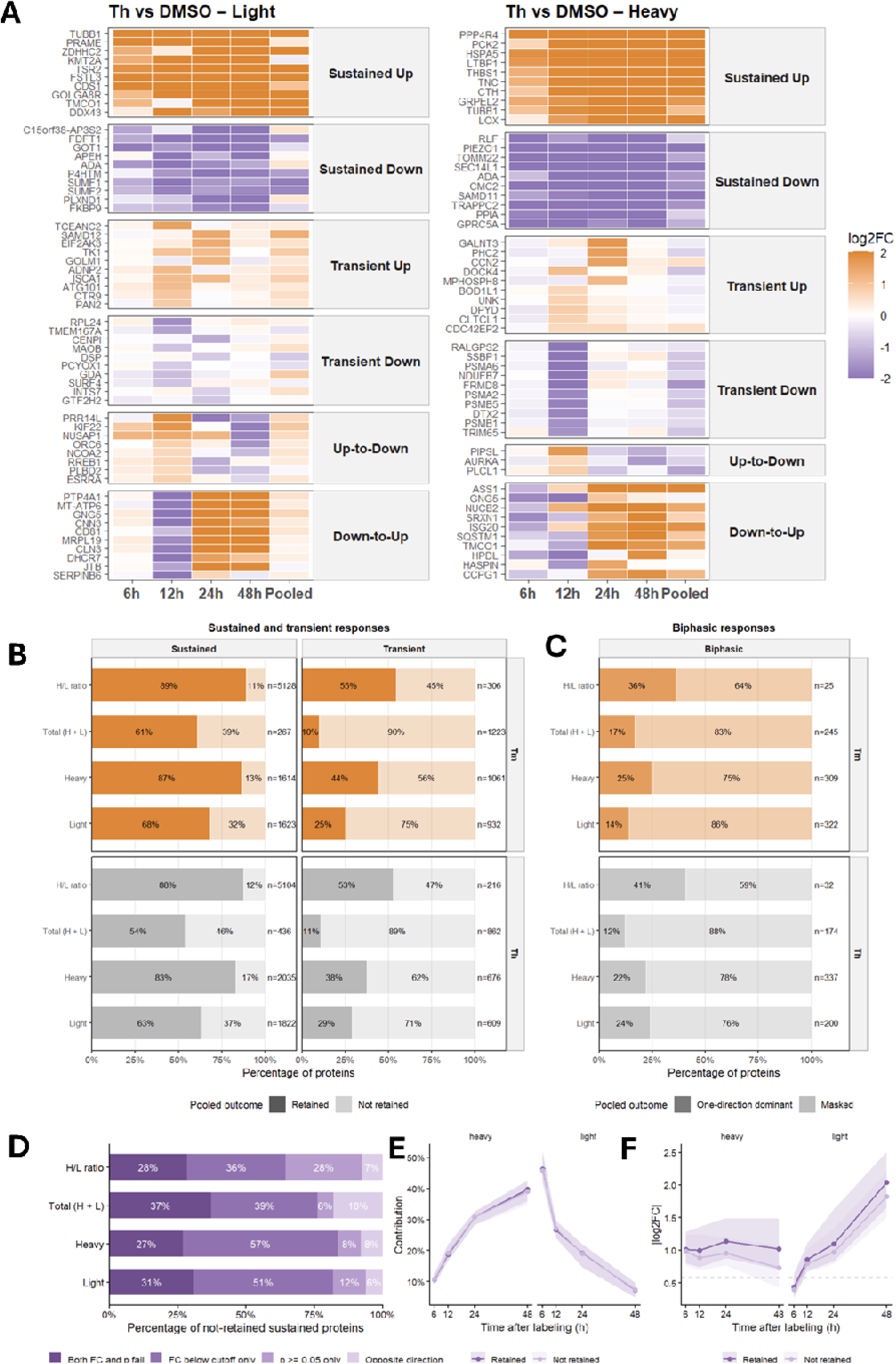
ISDia captures sustained and trainset responses but masks biphasic responses. **A.** Representative time-resolved patterns in light and heavy channels following Th treatment relative to DMSOalongside the corresponding ISDia pooled measurement. At each time point, proteins were defined as significantly increased (Up) or decreased (Down) using |log2FC| > 0.58 and p < 0.05 as cutoffs. Three categories of temporal protein responses are shown: Sustained: significant regulation in the same direction at ≥3 of 4 time points, with no significant regulation at the remaining time points. Transient: a temporally restricted increase or decrease that emerged and resolved within the time window, with no significant regulation at the remaining time points. Biphasic: significant changes in both directions, classified as Up-to-Down or Down-to-Up by response order. Up to 10 representative proteins with the most distinct temporal patterns are shown per response class. **B.** Bar plots showing the proportions of sustained and transient responses retained in the corresponding pooled ISDia measurements. Responses were classified as “Retained” when the pooled ISDia measurement detected significant regulation in the same direction, and as “Not retained” when the protein was no longer significantly regulated in the same direction after pooling. Bars show the percentage of proteins retained in each class. Numbers within bars indicate percentages, and n indicates the number of assessable proteins used as the denominator for each bar. **C.** Bar plots showing the retention rate of biphasic responses by ISDia pooled measurement. the responses were considered “Masked” when no significant change was detected in ISDia. When the pooled analysis detected a significant change, the response was classified as “One-Direction dominant”, indicating that one of the biphasic patterns predominated after pooling. **D.** Classification of sustained-response proteins not retained in ISDia based on fold-change and P-value thresholds. Percentages were calculated relative to the total number of non-retained sustained proteins in the Th-versus-DMSO comparison. **E.** Relative contribution of individual time points to pooled heavy and light intensity for retained and non-retained sustained response proteins in DMSO samples. For each protein, the contribution of a time point was calculated as the mean intensity at that time point divided by the sum of the mean intensities across all four time points (6, 12, 24, and 48 h). Lines and points represent the median contribution across proteins, and shaded regions indicate the interquartile range. **F.** Temporal response trajectories of retained and non-retained sustained proteins in the heavy and light channels under Th treatment relative to DSMO control. Lines and points show the median absolute log_2_ fold change at each time point, and shaded regions indicate the interquartile range. The dashed horizontal line indicates the fold-change cutoff of |log2FC| = 0.58. Retained proteins were those whose pooled responses remained directionally consistent and met both the fold-change and statistical significance criteria.

**Figure S5.**
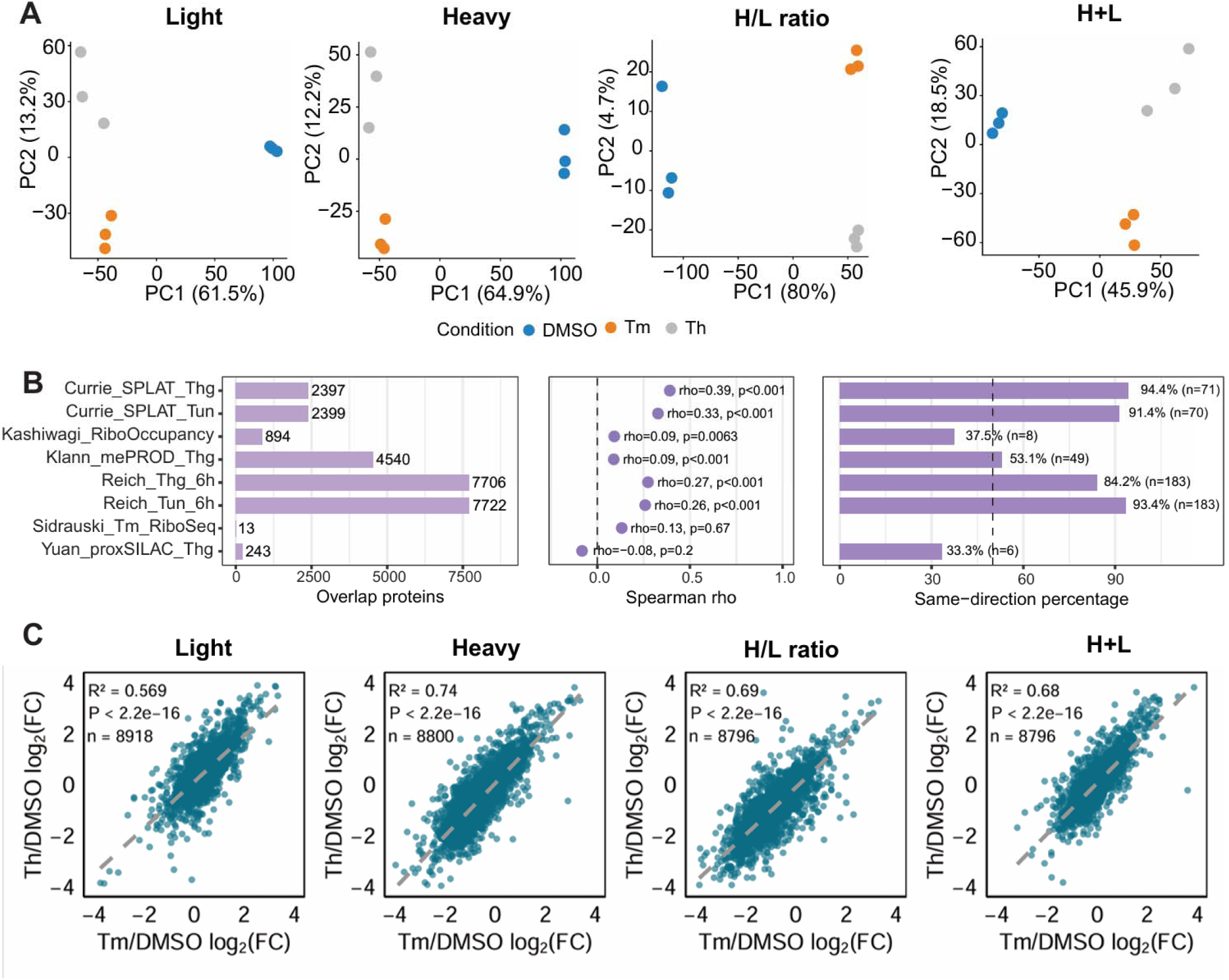
Quality control and validation of the ISDia dataset. A. Principal component analysis (PCA) of the WT dataset from three biological replicates for each treatment in ISDia. B. Comparison of ISDia results with previous studies. The left panel shows the number of overlapping genes/proteins between ISDia and each external dataset. The middle panel shows the Spearman correlation between ISDia and external datasets. Labels show Spearman’s rho, p-value, and the number of overlapping genes/proteins. The right panel shows the directional concordance between ISDia and external datasets among the most responsive matched gene products. With n each comparison, gene products were ranked by their effect values separately in the external dataset and in ISDia; the top 5% were labeled as upregulated and the bottom 5% as downregulated in each dataset. Only gene products classified as either upregulated or downregulated in both datasets were retained for analysis. Labels indicate the percentage of shared extreme gene products that changed in the same direction in ISDia and the corresponding external dataset, as well as the number of shared extreme gene products (n). C. Correlation between log_2_(Th/DMSO) and log_2_(Tm/DMSO) for heavy, light, and H+L intensities, as well as H/L ratio. n indicates the number of proteins included. R is the Pearson correlation coefficient, and p is the corresponding p-value.

**Figure S6.**
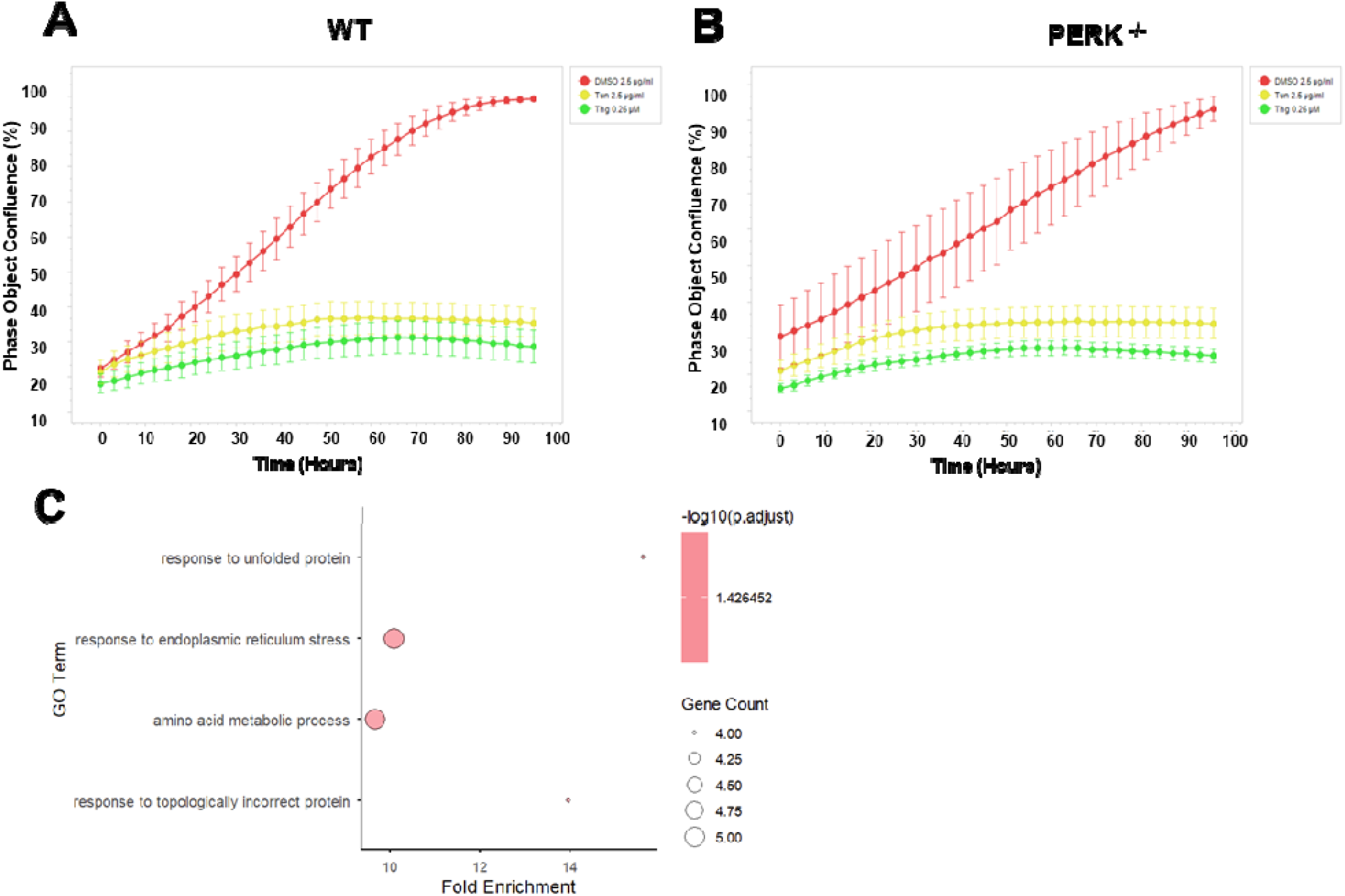
Adjusting for cell-division effects under UPR. A. Effects of ER stress on WT cell proliferation. Phase-object confluence was monitored over 96 h in cells treated with DMSO, tunicamycin (Tun), or thapsigargin (Thg). Confluence was measured from phase-contrast images and expressed as a percentage of the imaged area. Points represent the mean confluence at each time point, and error bars indicate variability among replicate wells. B. Effects of ER stress treatments on PERK ^-/-^ cell proliferation. Confluence was measured and calculated using the same method described in A. C. Biological processes enriched among the proteins in group A (Fig. 4A-B dark pink), defined as proteins with a log_2_ H/L(FC) > 0 under Th treatment relative to DMSO, based on light-channel values adjusted for cell division.

**Figure S7.**
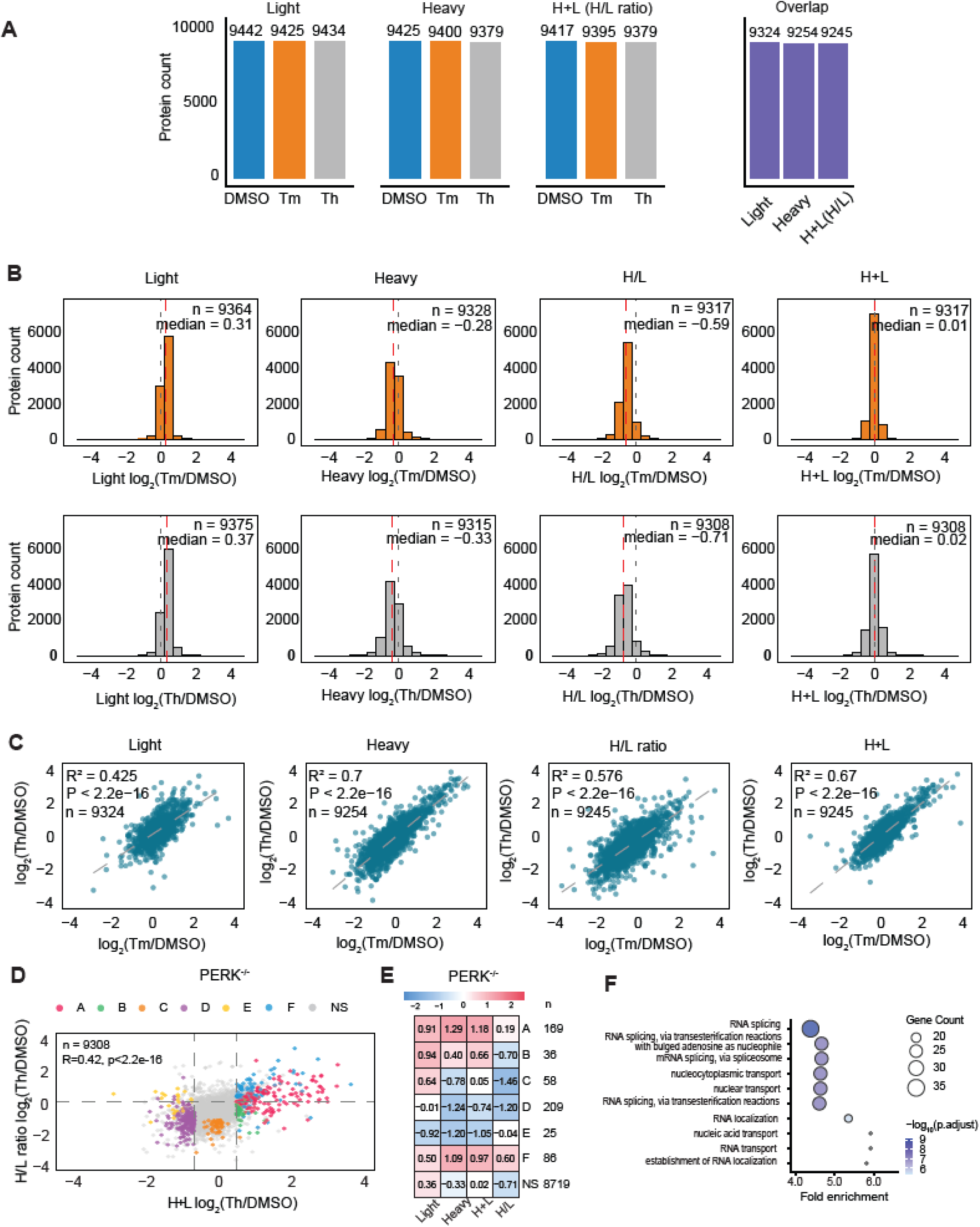
ISDia analysis of protein synthesis and clearance regulation in PERK^-/-^ cells during the UPR, related to Figure 4. A. Number of proteins quantified with light, heavy, and H+L (H/L ratio) across three replicates in each condition in PERK^-/-^ cells. H+L and H/L ratio values are included only when the corresponding l ght and heavy intensities were quantified in all three replicates. B. Distribution of log_2_ fold changes for light, heavy, H+L intensities and H/L ratios in PERK^-/-^ cells under Tm (top) and Th (bottom) treatment relative to DMSO. H+L and H/L ratio were included only when both light and heavy intensity were quantified in all three replicates. n indicates the number of included in each panel. Proteins C. Pairwise comparison of log_2_(Th/DMSO) and log_2_(Tm/DMSO) for light, heavy, H+L intensities and H/L ratios in PERK^-/-^ cells. n indicates the number of proteins included in each panel; R^2^ is the Pearson correlation coefficient; p is the corresponding p-value. D. Pairwise comparison of changes in H+L intensity and H/L ratio between Th and DMSO treatments in PERK^-/-^ cells. Protein labeling is the same as in Figure 4A. E. Heatmap showing the median log_2_ fold change in light, heavy, H+L intensity, and H/L ratio for each group in PERK^-/-^ cells. Values in each cell represent the median, and the color scale represents the magnitude and direction of the change (blue, decreased; red, increased). F. Biological processes enriched among proteins that showed decreased synthesis and decreased clearance in WT cells (down in heavy and up in light but H+L not significant), but no decreased synthesis in PERK^-/-^ cells under Th treatment compared to DMSO.

**Figure S8.**
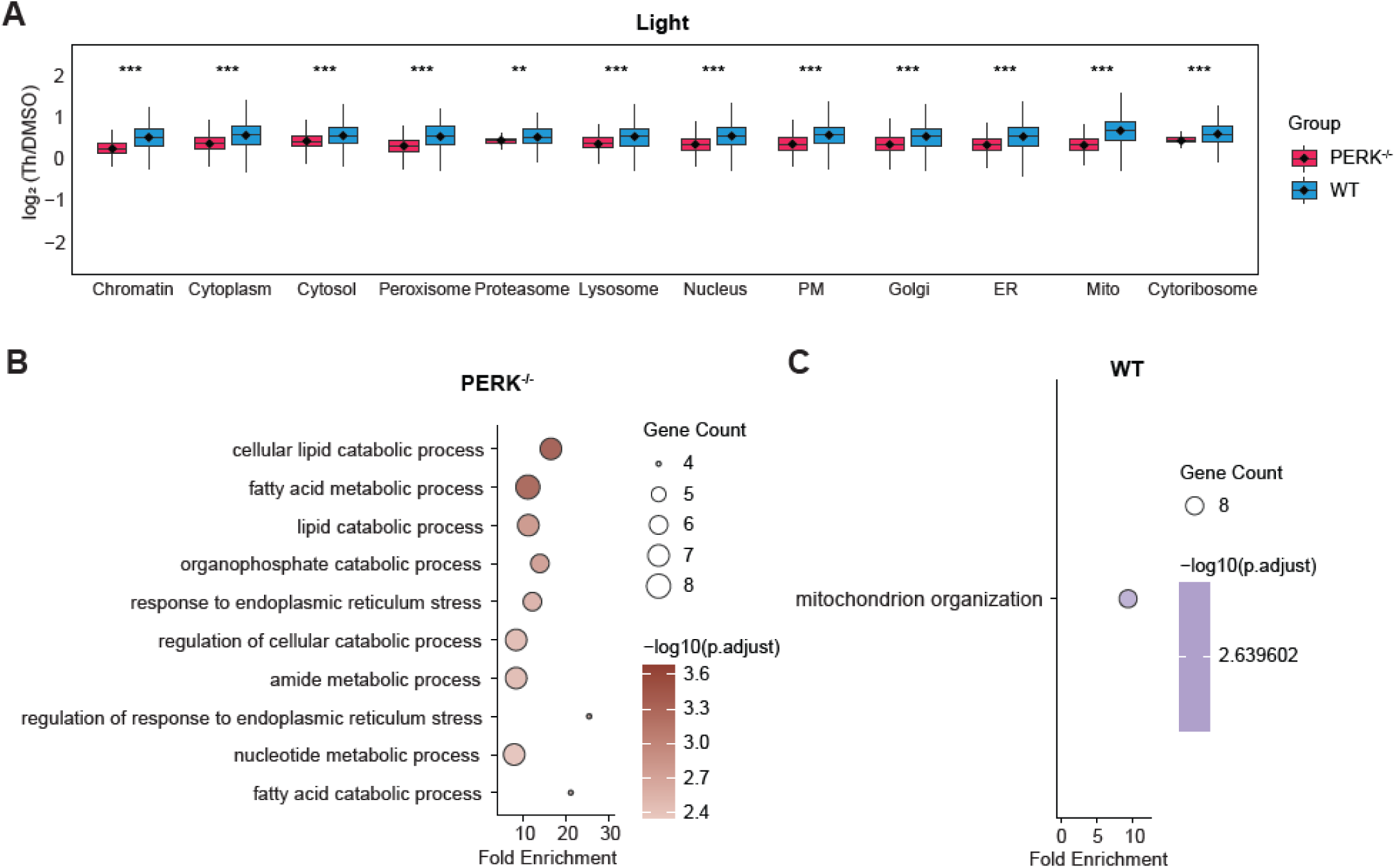
Regulation of mitochondrial protein synthesis and clearance during the UPR, related to Figure 5. A. Boxplot of log_₂_(Th/DMSO) fold changes for light intensities comparing WT and PERK^-/-^ cells across subcellular compartments. P values are from paired, one-tailed t-tests. Statistical significance is indicated as: *** p□<□0.001; **□0.001□<□p□<□0.01; *□0.01□<□p□<□0.05, ns, not significant. B. Biological processes enriched among mitochondrion (GO:0005739) proteins with significantly increased H+L intensity in PERK^-/-^ cells under Th treatment compared to DMSO (H+L log_2_(Th/DMSO) > 0.58 & p-value < 0.05). C. Biological processes enriched among mitochondrion (GO:0005739) proteins with significantly decreased H+L intensity in WT cells under Th treatment compared to DMSO (H+L log_2_(Th/DMSO) < - 0.58 & p-value < 0.05).

**Figure S9.**
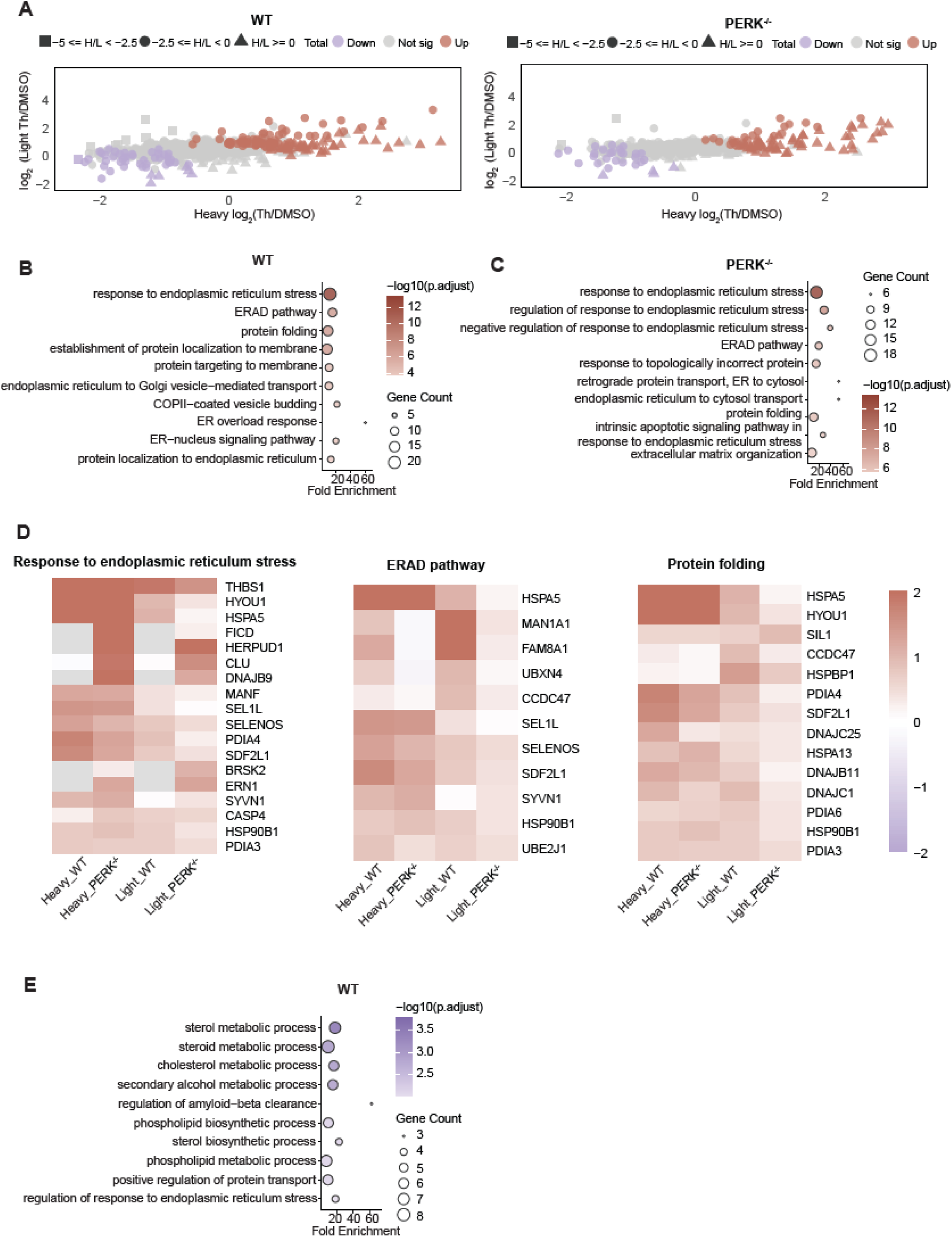
Regulation of ER protein synthesis and clearance during the UPR, related to Figure 5. A. Bubble plots showing log_₂_(Th/DMSO) changes in light, heavy intensity, H+L and H/L ratios for ER proteins ER in WT and PERK^-/-^ cells. B. Biological processes enriched among ER (GO:0005783) proteins showing significantly increased H+L intensity in WT cells under Th treatment relative to DMSO control (H+L log2(Th/DMSO) > 0.58 & p-value < 0.05). C. Biological processes enriched among ER (GO:0005783) proteins showing significantly increased H+L intensity in PERK^-/-^ cells under Th treatment relative to DMSO control (H+L log2(Th/DMSO) > 0.58 & p-value < 0.05). D. Heatmap of heavy and light intensity changes for proteins shown in **Fig. S5B** that are annotated to the following categories: response to endoplasmic reticulum stress (GO:0034976), ERAD pathway (GO:0036503) and protein folding (GO: 0006457) categories in WT and PERK^-/-^ cells.

**Figure S10.**
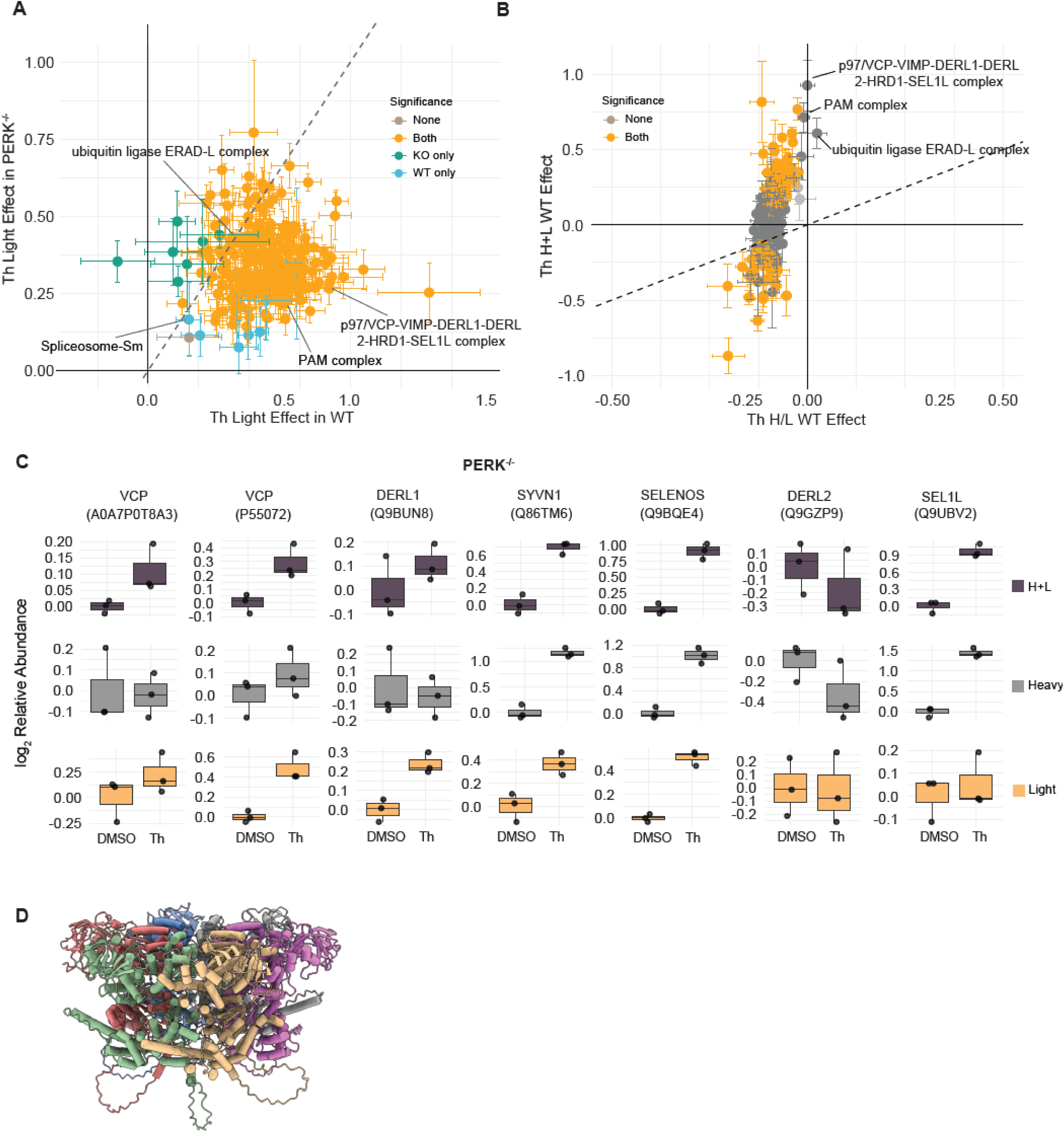
Differential regulation of synthesis and clearance among subunits across and protein complexes during the UPR, related to Figure 6. **(A)** Scatter plot showing H+L intensities and H/L ratios changes for protein complexes in WT cells under Th treatment relative to DMSO. Significance was indicated based on p-value < 0.05. Statistical significance was defined as an adjusted p-value < 0.05. Error bars represent the standard error. **(B)** Scatter plot showing changes in light intensities for protein complex subunit under Th treatment relative to DMSO in WT and PERK^-/-^ cells. Statistical significance was defined as an adjusted p-value < 0.05. Error bars represent the standard error. **(C)** Boxplot showing changes in H+L, heavy and light intensities subunits of the p97/VCP-VIMP-DERL1-DERL2-HRD1-SEL1L complex in PERK^-/-^ cells under Th treatment relative to DMSO. **(D)** Side view of the AlphaFold3-predicted hexameric structure of the short P97/VCP isoform (UniProt ID: A0AP0T8A3), with each protomer uniquely colored for clarity.

**Figure S11.**
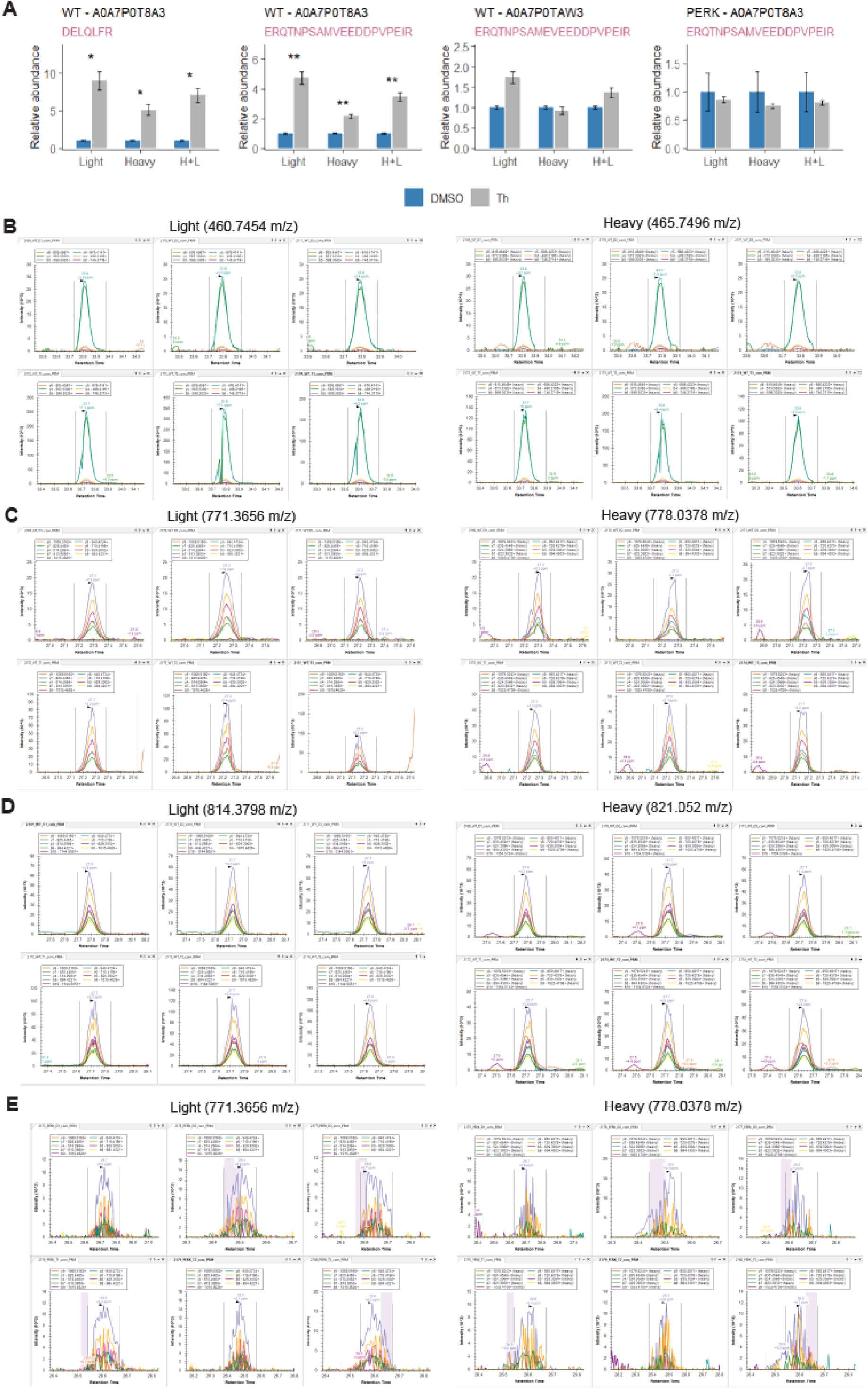
PRM validation of p97/VCP isoform expression changes quantified by ISDia. A. Bar plot showing the relative abundance of long- and short-isoform p97/VCP isoform-specific peptides under DMSO and Th treatments in WT and PERK^-/-^ cells, quantified by PRM using total fragment ion area. B. Representative light (left) and heavy (right) MS2 PRM chromatographic peaks for DELQLFR (2+ precursor) in WT cells. C. Representative light (left) and heavy (right) MS2 PRM chromatographic peaks for ERQTNPSAMVEEDDPVPEIR (3+ precursor) in WT cells. D. Representative light (left) and heavy (right) MS2 PRM chromatographic peaks for ERQTNPSAMEVEEDDPVPEIR (3+ precursor) in WT cells. E. Representative light (left) and heavy (right) MS2 PRM chromatographic peaks for ERQTNPSAMVEEDDPVPEIR (3+ precursor) in PERK^-/-^ cells.

**Figure S12.**
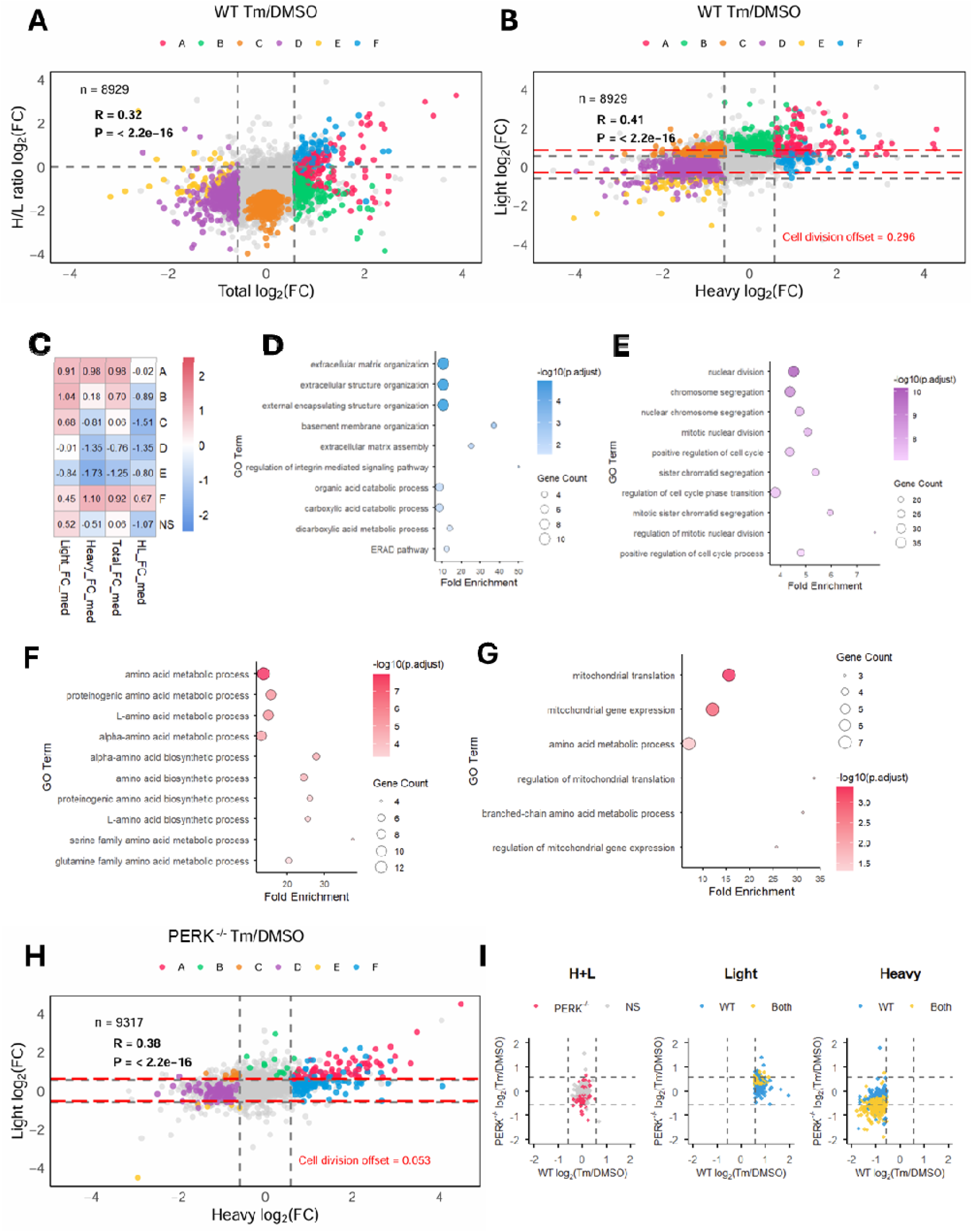
Diverse protein synthesis and clearance regulation shapes proteome abundance during the UPR triggered by Tm treatment. A. Scatter plot showing changes in H+L and H/L ratio between Tm and DMSO treatments. Each point represents a protein. Grey dashed lines indicate the fold change thresholds of log_₂_(1.5) ≈ ±0.58 for H+L intensity, and log_₂_(1) = 0 for H/L ratio. Proteins are color-coded based on their changes across light, heavy, and H+L as follows: A: dark pink (up in light, heavy, and H+L); B: green (up in light and H+L, heavy not significant); C: orange (light up, heavy down, H+L not significant); D: purple (down in heavy and H+L, light not significant); E: yellow (down in light, heavy, and H+L); F: blue (up in heavy and H+L, light not significant) grey (others). Up is log_2_FC > 0.58 & p-value < 0.05; Down is log_2_FC < -0.58 & p-value < 0.05; no significant is p-value >0.05. n is the number of proteins shown; R is the Pearson correlation coefficient; p is the corresponding p-value. B. Scatter plot showing changes in heavy and light intensities between Tm and DMSO treatment. Color scheme and significance thresholds are the same as in A. Red dashed lines indicate the cell division-adjusted light-channel fold-change thresholds, calculated as −0.58 + the cell division offset and +0.58 + the cell division offset. Cell division offset in WT is 0.296. C. Heatmap showing the median log_2_ fold change in light, heavy, H+L intensity, and H/L ratio for all proteins within groups highlighted in A and B. Values in each cell represent the group-wise median. D. Biological processes enriched among proteins in group F (blue). E. Biological processes enriched among proteins in group D (purple). F. Biological processes enriched among the proteins in group A (dark pink) with log2 H/L(FC) > 0 under Tm treatment compared to DMSO. G. Biological processes enriched among the proteins in group A (dark pink) with log2 H/L(FC) < 0 under Tm treatment compared to DMSO. H. Scatter plot showing the changes in H+L and H/L ratio between Tm and DMSO treatment in PERK^-/-^ cells. Color scheme and significance thresholds are the same as A. Red dashed lines indicate the cell division-adjusted light-channel fold-change thresholds, calculated as −0.58 + the cell division offset and +0.58 + the cell division offset. I. Scatter plot showing the log_2_(FC) in heavy, light and H+L intensities under Tm treatment relative to DMSO between WT and PERK^-/-^ cells for group C (orange group) proteins filtered from WT cells (light up, heavy down, H+L not significant). Each point represents one protein from this defined group in WT, plotted by its corresponding log_2_(fold change) under Tm relative to DMSO in WT and PERK^-/-^ cells. Colors represent significance of the displayed metric: gray, non-significant in both cell types; pink, significant only in PERK^-/-^ cells; blue, significant only in WT cells; yellow, significant in both cell types. Statistical significance: p□<□0.05.

**Figure S13.**
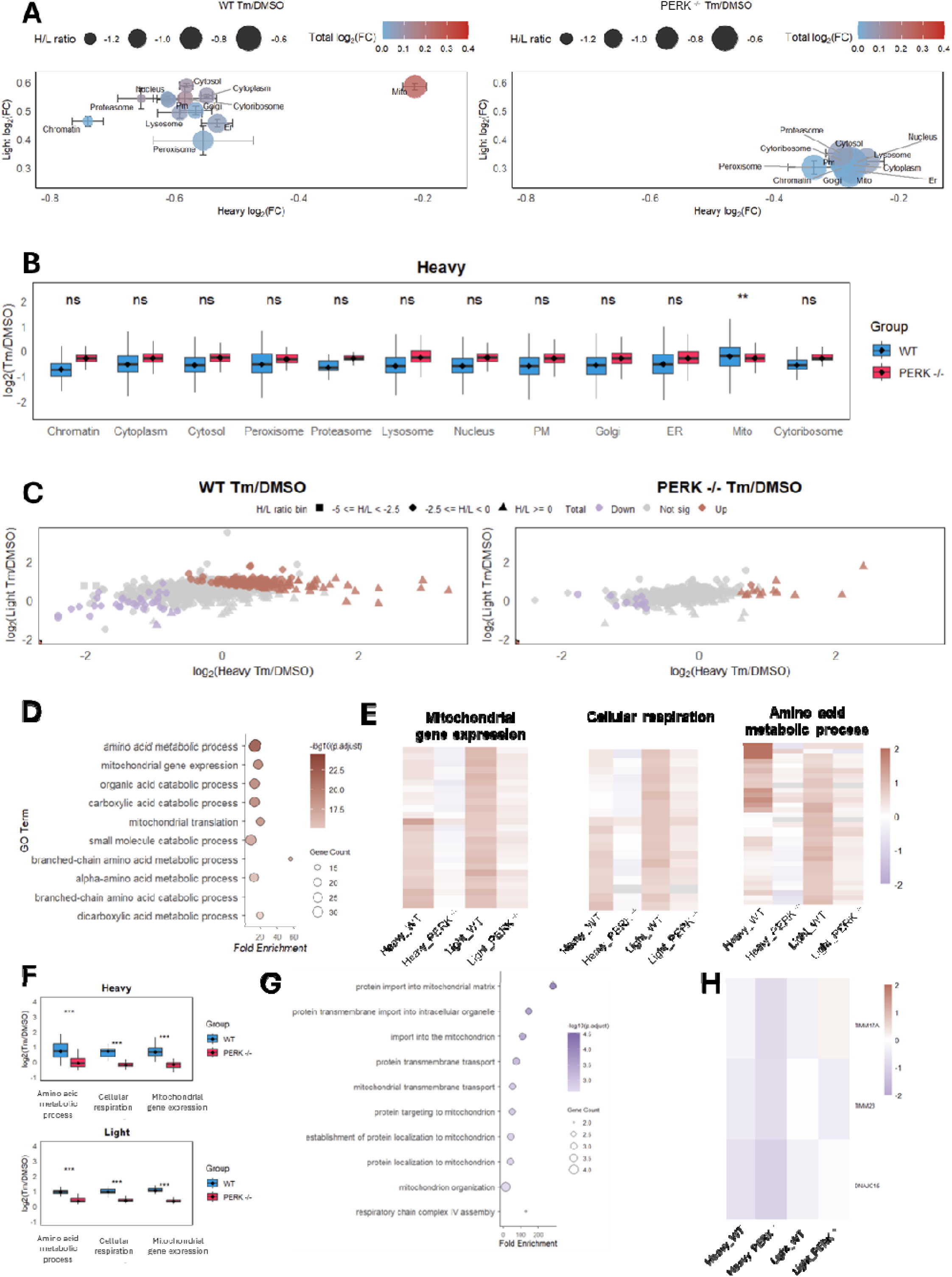
Differential regulation of protein synthesis and clearance across cellular compartments in WT and PERK^-/-^ cells under Tm treatment. A. Bubble plots showing the median of log2(FC) in light, heavy, H+L, and H/L ratio across subcellular compartments in WT (left) and PERK^-/-^ (right) cells under Tm treatment relative to DMSO. Error bars represent the standard deviation. B. Boxplot of log_₂_(FC) in heavy peptide intensities across subcellular compartments in WT and PERK^-/-^ cells under Tm treatment. P values are from paired, one-tailed t-tests. ***□p□<□0.001; **□0.001□<□p□<□0.01; *□0.01□<□p□<□0.05; ns, p□>□0.05. C. Bubble plots showing log_₂_(FC) in light, heavy, H+L, and H/L ratios, for mitochondrial proteins in WT and PERK^-/-^ cells following Tm treatment relative to DMSO. D. Biological processes enriched among mitochondrial proteins with significantly increased total abundance in WT cells following Tm treatment relative to DMSO (H+L log_2_(Th/DMSO) > 0.58 & p-value < 0.05). E. Heatmap of heavy and light intensity changes for proteins belonging to selected enriched mitochondrial categories from D, including mitochondrial gene expression (GO:0140053), cellular respiration (GO:0045333), and amino acid metabolic process (GO:0006520). F. Boxplot of log_₂_(Tm/DMSO) fold changes in heavy and light peptide intensities comparing WT and PERK-/- cells across categories in E. P values were calculated using paired, one-tailed t-tests. ***□p□<□0.001; **□0.001□<□p□<□0.01; *□0.01□<□p□<□0.05; ns, not significant. G. Biological processes enriched among the mitochondrial proteins with significantly decreased total abundance in PERK^-/-^ cells (H+L log_2_(Tm/DMSO) < -0.58 & p-value < 0.05). H. Heatmap of heavy and light intensity changes for proteins annotated to the enriched protein targeting to mitochondrion (GO:0006626) category from G in WT and PERK^-/-^ cells.

**Figure S14.**
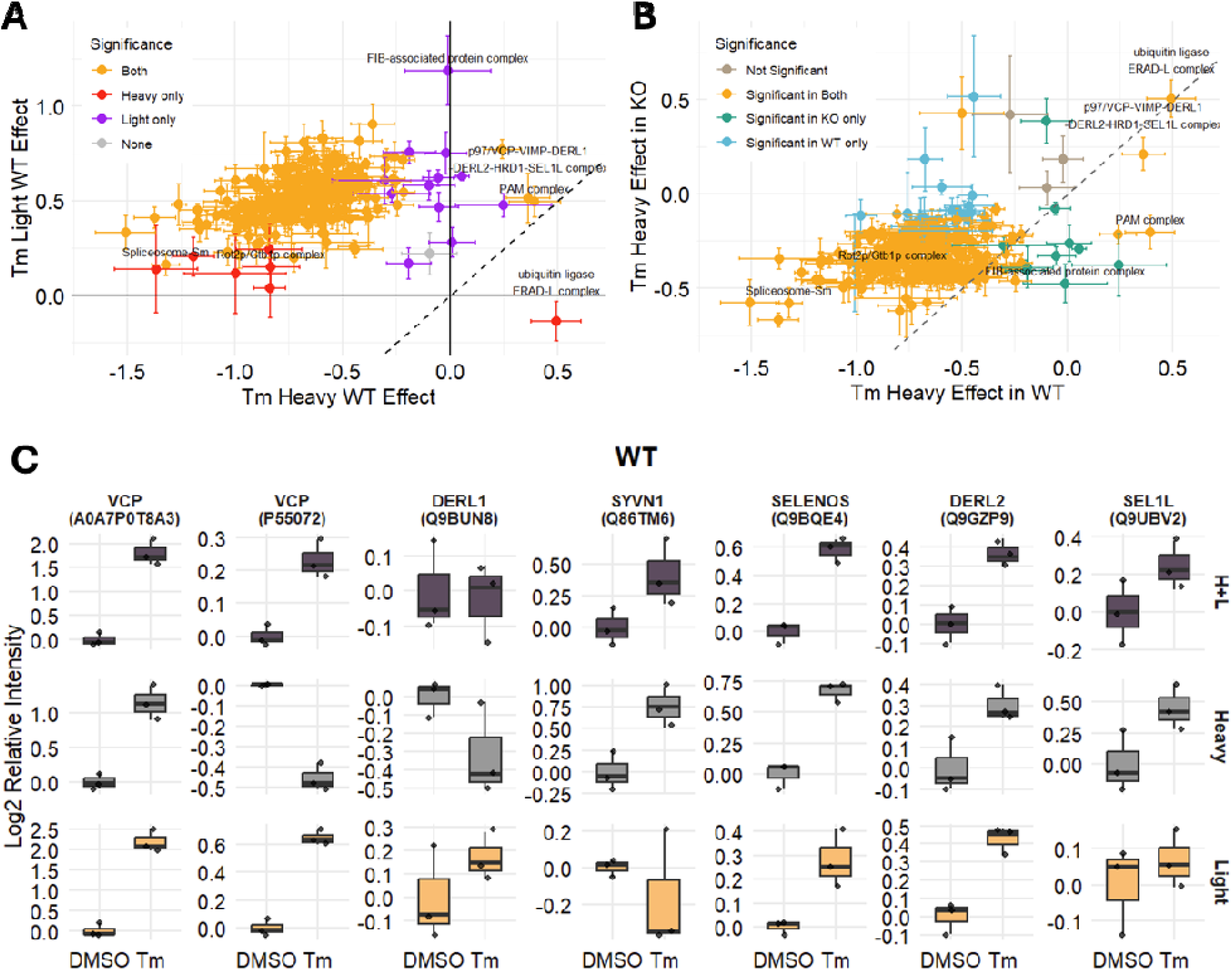
Differential regulation of synthesis and clearance among subunits across and within protein complexes under Tm treatment. A. Scatter plot showing the changes in heavy and light intensities for protein complexes in WT cells treated with Tm relative to DMSO. Statistical significance was defined as an adjusted p-value < 0.05. Error bars represent the standard error. B. Scatter plot showing the changes in heavy peptide intensities for protein complexes under Tm treatment relative to DMSO in WT and PERK-/- cells. Statistical significance was defined as an adjusted p-value < 0.05. Error bars represent the standard error. C. Boxplots showing changes in heavy, light and H+L intensities for subunits of p97/VCP-VIMP-DERL1-DERL2-HRD1-SEL1L complex in WT cells under Tm treatment relative to DMSO.

## Supplemental Table

**Table S1**. Changes in heavy and light peptide intensities, as well as H + L abundance and H/L ratios, at each time point following Th or Tm treatment in WT cells. Related to **Figure 2**.

**Table S2**. Changes in heavy and light peptide intensities, as well as H+L and H/L ratios from ISDia, under Th or Tm treatment in WT cells. Cellular compartments are annotated. Related to **Figure 3-5**.

**Table S3**. Summary of external datasets used for comparison with ISDia. Related to **Figure S5B**.

**Table S4**. Changes in heavy and light peptide intensities, as well as H+L and H/L ratios from ISDia, under Th or Tm treatment in PERK^-/-^ cells. Cellular compartments are annotated. Related to **Figure 4-5**.

**Table S5**. Changes in heavy and light peptide intensities, as well as H+L and H/L ratios of protein complexes, under Th or Tm treatment in WT and PERK^-/-^ cells. Related to **Figure 6**.

**Table S6.** Inclusion table of PRM. Related to **Figure S11**.

